# Resilience of rapeseed to heat stress during grain filling in a high yielding environment

**DOI:** 10.1101/2024.12.11.628060

**Authors:** José Verdejo, Daniel F. Calderini

## Abstract

Global climate change is driving the temperature increase, which negatively impacts on crop production. Most heat stress studies in rapeseed have been conducted under controlled conditions, limiting their results to true field crops. This study aimed to assess the sensitivity of rapeseed to temperature increase during two phases of the grain filling period in southern Chile, a high-yield potential environment. To our knowledge, this is the first field study evaluating the effects of heat stress at different phases of grain filling in rapeseed. Three field experiments were conducted with two adapted spring rapeseed hybrids, Lumen and Solar CL, under three temperature treatments: a control at ambient temperature, a 5°C increase from the beginning of flowering to 15 days after flowering (DAF), and the same increase from 15 to 30 DAF. Crop and climate variables, including air temperature and solar radiation were recorded along the experiments. Slight effects on grain yield due to heat stress were found, however, the hybrids exhibited different sensitivities, with Lumen being less affected than Solar CL. Grian yield of the last hybrid showed positive association with photothermal quotient and negatively with temperature, while Lumen did not show relationship. The most significant impact on grain yield occurred during the first half of grain filling (0-15 DAF), resulting in a reduction of 26.8% of grain number in Solar CL, compared to a 6.0% reduction in Lumen across experiments. Grain weight remained little affected by thermal stress, indicating its conservative behaviour and tolerance in southern Chile conditions. Grain oil concentration was scarcely sensitive, while grain protein concentration increased under heat stress. The low impact on the crop outcomes of our studies may be attributed to the lower background temperature of southern Chile, suggesting that this environment may confer greater rapeseed heat tolerance during grain filling than in other agroecosystems.

## 1. Introduction

Unavoidable global climate change involves increases in atmospheric CO_2_ concentration, average temperature, and rainfall variation, along with a higher frequency of extreme climate events (Asseng et al., 2015; Lee et al., 2021). Global warming of 1.5°C implies higher mean temperatures compared to the 1850–1900 period, with generally more significant warming over land than over ocean areas and larger increases in high latitudes compared to low latitudes (Lee et al., 2021). The rise in temperature due to climate change is expected to negatively impact on crop production (e.g. Porter and Semenov, 2005; Ray et al., 2019; Trnka et al., 2004) as relatively small changes in average temperature have shown significant effects on crop productivity (Cheng et al., 2009; Kim and You, 2010). Although there is agreement on the forecasted impact of global warming on crops, current knowledge about crop adaptation to heat conditions remains incomplete.

In the Southern Hemisphere, the strongest warming over land is expected to occur in the subtropical areas of South America, Southern Africa, and Australia, regardless of the level of global warming (Lee et al., 2021). Temperature increases have already been observed in the Southern Cone of South America over the past 50 years (Rivelli et al., 2021). If this trend continues, global warming will endanger the current adaptation of crops, considering that temperature is a key environmental factor regulating both the development rate and growth of crops (Egli, 2017; Sadras and Calderini, 2021; Slafer and Rawson, 1994). In light of this, agriculture in southern Chile, one of the world’s highest yield potential areas for temperate crops, will be negatively affected, as has already been reported for various crops (Lesjak and Calderini, 2017; Lizana and Calderini, 2013; Mera et al., 2015; Sandaña et al., 2009; Verdejo and Calderini, 2020). Therefore, rapeseed production, which is widely grown in this area, is challenged by global warming, especially during the grain filling period, according to the scenarios of increased temperatures forecasted for southern Chile (Araya-Osses et al., 2020; Hernandez and Madeira, 2022).

Many studies have evaluated responses to increased temperatures in cereals, grain legumes, and also oilseed crops (e.g., Awasthi et al., 2014; Egli et al., 2005; García et al., 2015; Liu et al., 2022; Lizana and Calderini, 2013; Mayer et al., 2014; Rattalino Edreira et al., 2011; Rondanini et al., 2006; Sehgal et al., 2017). However, less effort has been dedicated to the impact of higher temperatures on rapeseed, the third most consumed plant oil after palm and soybean (UFOP, 2024), which produces one of the healthiest plant oils for human consumption (Konuskan et al., 2019). Previous heat stress evaluations of this crop showed grain yield reductions of 30% (Pokharel et al., 2021; Triboi-Blondel and Renard, 1999), due to decreases in grain number (17-40%) and grain weight (19-40%). In addition to rapeseed productivity, the impact of heat on quality traits has shown oil reductions of 12% and a trade-off with protein concentration, whereby this last trait increased from 16 to 38% in response to raised temperatures (Pokharel et al., 2021; Triboi-Blondel and Renard, 1999). However, current heat stress studies in rapeseed were mainly carried out under controlled conditions, preventing simple extrapolations to field conditions (Secchi et al., 2023). Often, controlled environments use unrealistic approaches since they include the use of plants in small soil volume containers, lacking natural radiation and having modified vapour pressure deficits (Bonada and Sadras, 2015). In this regard, it is important to take into account that the sensitivity of both grain yield and quality traits of crops depends on the degree of temperature increase, the duration of this increase, the developmental phase of the crop, the ambient background temperature, and interactions among these factors and with other variables including genotype and management practices (Cossani and Sadras, 2021; Sadras et al., 2022; Sadras et al., 2013; Slafer et al., 2014). Therefore, heat stress evaluations of rapeseed under field conditions are needed to develop a precise cause-and-effect framework for the crop responses to temperature increases.

In light of the global warming scenarios for southern Chile, the temperature increase will mainly affect the post-flowering period (from BBCH55 onwards) of rapeseed. However, few experiments have been conducted on this crop to assess the temperature increase during grain filling, especially under field conditions (Pokharel et al., 2021; Rivelli et al., 2024; Rivelli et al., 2023). Additionally, the effect of increased temperature during different phases of grain filling has been assessed in other crops (e.g., sunflower: Rondanini et al. (2006); wheat: Lizana and Calderini (2013); Stone and Nicolas (1995); maize: Wang et al. (2022); soybean: Soba et al. (2022) and references therein). But, to the best of our knowledge, there are no previous studies on rapeseed reporting the impact of heat stress at different phases of grain filling in field experiments.

Like other grain crops, rapeseed has demonstrated a critical period for grain number determination (Kirkegaard et al., 2018). This period spans from 100 to 500 cumulative degree days (°Cd) after the start of flowering. In line with this, Verdejo and Calderini (2020) found that the sensitivity of this crop to reductions in the source-sink ratio was greater between 0 and 15 days after flowering (DAF) than from 15 to 30 DAF in spring rapeseed. Moreover, we reported contrasting resilience in grain yield between the first and second periods of grain filling, attributed to the ability of grain weight to increase in response to reductions in grain number under source limitations from 0 to 15 DAF. The positive response of rapeseed grain weight, allowing for full or partial compensation to reductions in grain number, has also been noted in other studies on this crop (Kirkegaard et al., 2018; Labra et al., 2017; Rivelli et al., 2024). Therefore, the sensitivity of rapeseed grain weight, and consequently grain yield, to increased temperatures may vary significantly depending on the timing of thermal stress during grain filling. Furthermore, the considerable resilience exhibited by rapeseed grain weight has been scarcely evaluated under heat stress.

This background raises the question of whether the response of grain weight, grain yield, and quality traits (grain oil and protein concentrations) exhibited by rapeseed to source-sink reduction (Rivelli et al., 2024; Verdejo and Calderini, 2020) could be similar under increased temperatures during grain filling, considering that the physiological mechanisms involved in the response to these stresses are different (Campbell, 1981; Lambers and Oliveira, 2019; Rivelli et al., 2023; Zhang et al., 2022). The present study aims to assess the sensitivity of rapeseed grain yield, its numerical components, and grain quality traits, along with their physiological bases, to elevated temperatures during two-time windows (0-15 and 15-30 DAF) in the high-yield potential environment of southern Chile.

## 2. Materials and Methods

### 2.1. Experimental set-up

Three field experiments were performed at the Austral Farming Experimental Station (EEAA) of the Universidad Austral de Chile in Valdivia, Chile (39° 47’ S, 73° 14’ W), on a Duric Hapludand soil. In each experiment, two adapted spring rapeseed hybrids (Lumen and Solar CL from NPZ-Lembke, Germany) were assessed under three temperature treatments: (i) a control at ambient temperature, (ii) a 5°C temperature increase from the beginning of flowering [BBCH 61] to 15 days after flowering (DAF), and (iii) the same temperature increase from 15 to 30 DAF. Experiments 1 and 2 were sown on 5 September 2019 in different paddocks, while Experiment 3 was sown on 4 September 2020. Plots were arranged in a split-plot design with three replicates in Experiments 1 and 2, where the heat stress treatments were assigned to the main plots and genotypes to the sub-plots. In Experiment 3, treatments were arranged in a randomised block design with three replicates, where genotypes and heat stress treatments were fully randomised within each block.

### 2.2. Experimental management

Across the experiments, plots were 2 m long and 3.5 m wide, comprising 11 rows spaced 0.175 m apart. Each plot was sown at a seed rate of 55 plants m^-2^. At sowing, 150 kg ha⁻¹ of Ca(H_2_ PO_4_)_2_ and 100 kg ha⁻¹ of K_2_SO_4_·MgSO_4_ were applied to each experimental unit to prevent nutritional shortages of phosphorus, potassium, magnesium, and sulphur. No additional nitrogen was applied in Experiment 1, whereas in Experiments 2 and 3, nitrogen fertilisation was split into two applications of 100 kg N ha⁻¹ each, applied shortly after plant emergence (BBCH 10) and when the fifth internode expanded (BBCH 35). Nitrogen was applied in both experiments in the form of (NH_4_)NO_3_+ CaCO_3_+ MgCO_3_ (50% NO_3_ and 50% NH_4_).

Optimal management practices were implemented in the experiments. Plots were drip irrigated as required, depending on rainfall, until physiological maturity to prevent water shortages. Diseases, pests, and weeds were prevented or controlled in accordance with the rates recommended by the manufacturers to keep the experiments free of biotic constraints.

### 2.3. Heat stress treatments

The increase in temperature during the heat stress treatments was achieved using portable greenhouse chambers set up on the plots. The greenhouse structures were constructed with a wooden frame covered with 100 µm thick transparent polyethylene film on the treated plots (Fig. 1). The top of each structure was positioned 0.3-0.4 m above the plants during the treatments. The east side of the chambers was left open during daylight hours to allow for air circulation and pollinator access, similar to previous studies conducted at the EEAA (Labra et al., 2017; Verdejo and Calderini, 2020). Thermo-fans with a capacity of 2000 W were installed inside the greenhouses to raise the temperature by 5°C above ambient levels. Windows on the west side were opened when appropriate to prevent temperature increases beyond 5°C. Additionally, automatic sensors located inside and outside the greenhouses were employed to monitor this thermal difference. Thermal sensors also recorded temperature and relative humidity both inside and outside the greenhouses. To assess the effect of the polyethylene film on incident solar radiation reaching the crop, a linear ceptometer (AccuPAR LP-80, Decagon, Washington, USA) was utilised to measure radiation above and below the chambers once a week at midday during the heat stress treatments (four occasions). As a result of these measurements, the radiation intercepted by the polyethylene film averaged 10%.

**Figure 1.**
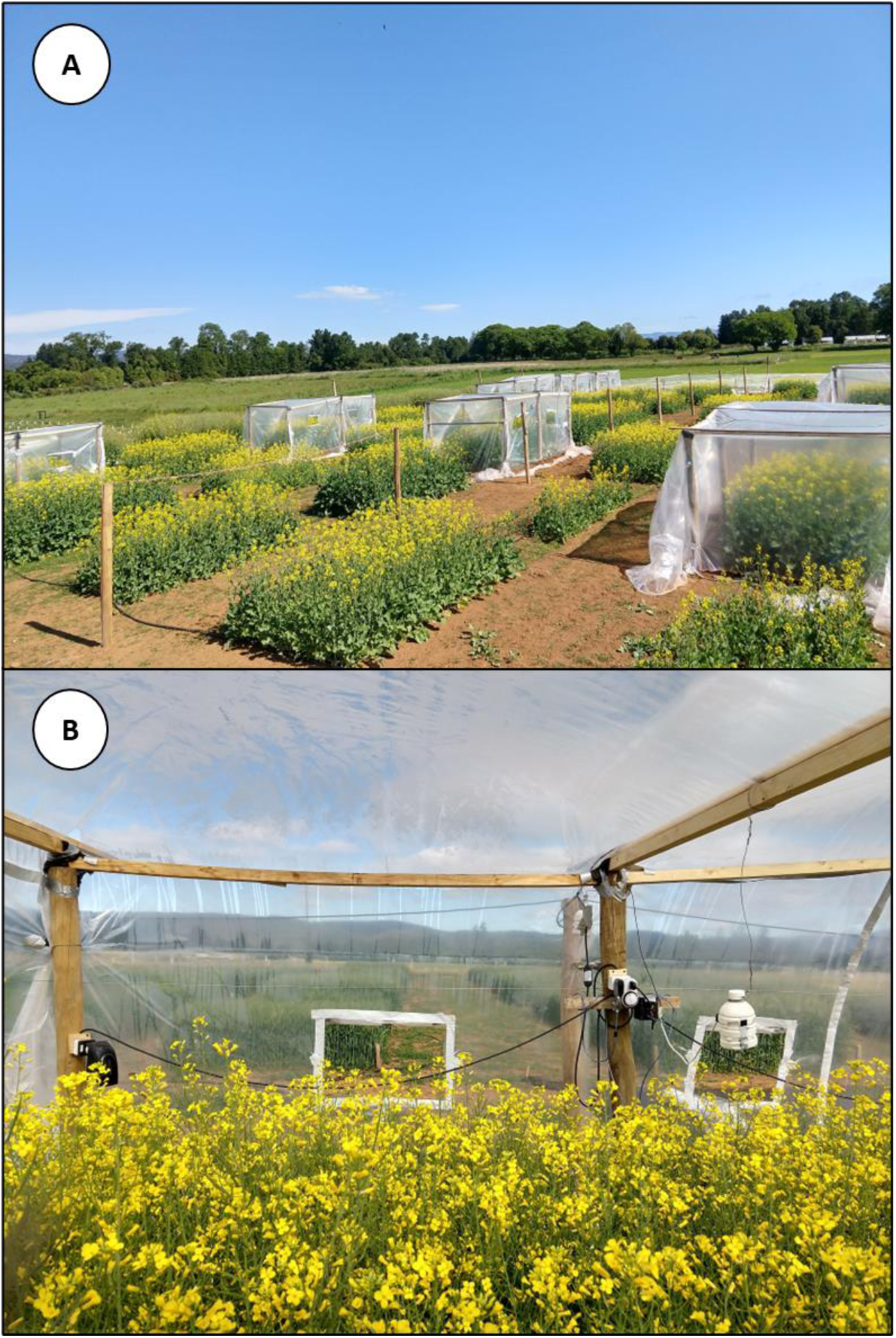
Experimental distribution of heat stress treatments during grain filling. (A) plastic greenhouses established on the plots and the internal layout of the greenhouse, (B) thermo-fans and the automatic sensors used to increase temperature.

### 2.4 Weather data

Maximum (Tmax) and minimum (Tmin) air temperatures and incident solar radiation (ISR) were recorded every 30 minutes from sowing until harvest at the Austral Meteorological Station of the EEAA (http://agromet.inia.cl/), located 200 m from the experiments. Thermal time (TT) was calculated by summing daily mean temperatures (Tmean) using a base temperature of 0°C (Kirkegaard et al., 2012). Heat load (temperature above 30°C) was calculated by summing hourly temperatures above 30°C throughout the days of the heat stress treatments, as described by Rivelli et al. (2024), based on the concept outlined by Wardlaw et al. (2002) and Rondanini et al. (2006). The photothermal quotient (PTQ) was calculated during the crop cycle as the ratio between daily ISR and Tmean. The effect of radiation reduction due to the polyethylene film (approximately 10%) was accounted for when applicable. PTQ was weighted by vapour pressure deficit (VPD) in accordance with Rodriguez and Sadras (2007). PTQ weighted by vapour pressure deficit (PTQ_VPD_) was calculated as the ratio between PTQ and VPD.

### 2.5. Crop measurements

Across the experiments, crop development was monitored twice a week according to the BBCH phenological scale for rapeseed (Meier, 2018). Rapeseed phenology was recorded at sowing, emergence, BBCH55, BBCH61 (start of flowering), 15 days after flowering (DAF), 30 DAF, and physiological maturity. The crop cycle and developmental phases were expressed in thermal time units, as mentioned above. At maturity, plant samples were harvested from one linear metre of the central rows in each plot of the experiments, when the grains inside the siliques were dark and hard (BBCH 89). Biomass samples were oven-dried at 65°C for 48 hours. Siliques from each plot sample were threshed and then weighed. Grain yield (GY), harvest index (HI), grain number (GN), number of siliques, weight per silique, grains per silique, thousand grain weight (TGW), and quality traits (grain oil and protein concentrations) were measured or calculated. Above-ground biomass, grain yield, and silique weight were measured using a precision balance (Radwag WTC 2000, Radom, Poland). Grain number was counted using a grain counter (Pfeuffer GmbH, Kitzengen, Germany). The harvest index was calculated as the ratio of grain yield to above-ground biomass. The weight per silique was calculated as the ratio of silique weight to number. The number of grains per silique was determined as the ratio of the total grain number per plant to the total number of siliques per plant. The average grain weight (thousand grain weight) was estimated as the ratio of grain yield to grain number. The oil concentration of grains was determined by Near Infrared Reflectometry (NIR) (Foss Infratec 1241, Hilleroed, Denmark), and the nitrogen concentration of grains was measured using the Kjeldahl procedure (Kirk, 1950). Protein concentration in grains was calculated using a conversion factor of 5.8 (Merrill and Watt, 1973). Concentrations of both oil and protein were expressed on a dry matter basis.

### 2.6. Measurement of individual grain dynamics

Fresh and dry weights of individual grains were measured throughout the grain filling period from the main raceme. These evaluations were conducted from the start of flowering to harvest maturity to account for the effects of heat stress on grain growth rate and grain filling duration. Three siliques on the main raceme (bottom, middle, and top positions) were sampled twice a week. Subsequently, the number of grains was counted, and the individual grain weight and water content were measured or calculated. Grains were oven-dried at 65°C for 48 hours and weighed using an analytical balance (Mettler, Toledo XP205DR, Greifensee, Switzerland). Individual grain water content was calculated as the difference between fresh and dry grain weight.

The time course of individual grain weight on the main raceme during grain filling was estimated using a tri-linear broken-stick function, as described by Verdejo and Calderini (2020). Individual grain weight was also assessed in relative terms, calculated as the measured grain weight relative to the final grain weight at physiological maturity. Relative individual grain weight was calculated for each treatment in each experiment. Additionally, the time course of individual grain water content in absolute terms was fitted using a centred third-order polynomial model for each treatment in all experiments, as detailed by Menendez et al. (2019). A centred model, which avoids computational problems such as overflows, was employed (Motulsky, 2014).

### 2.7. Statistical analysis

To assess the effectiveness of the heat stress treatments, the temperature was recorded both inside and outside the greenhouses once per hour. The daily average temperature difference recorded during the heat stress treatment period was compared using an extra-sum of squares F test in Statgraphics Centurion 18. Temperature changes throughout the heat stress treatment periods were plotted using GraphPad Prism 8 (Fig. 2).

**Figure 2.**
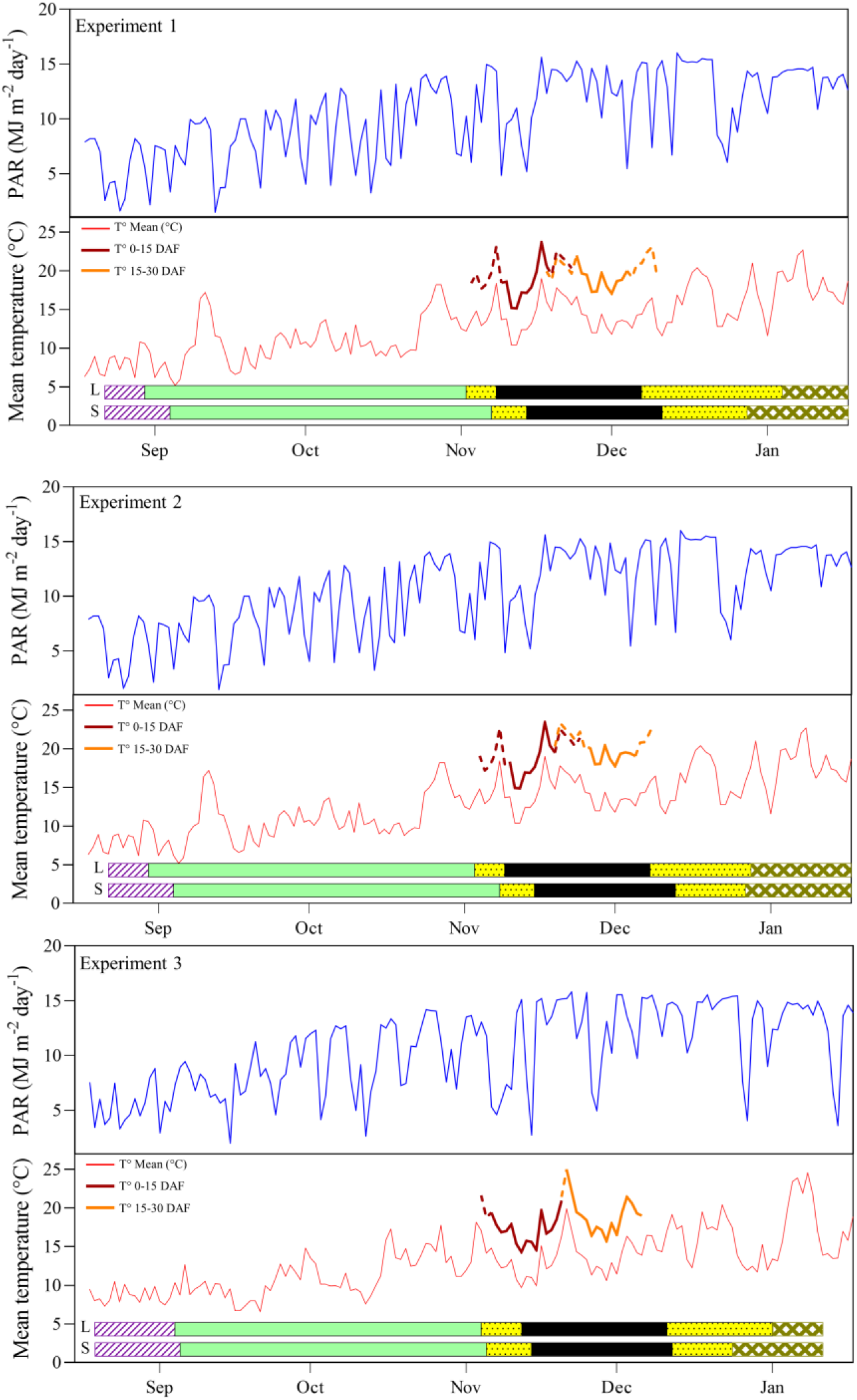
Photosynthetically active radiation (PAR), mean daily temperature and crop phenology of the genotypes (Lumen (L) and Solar CL (S)) in response to heat stress during grain filling across the experiments. The bars represent the crop phenology during the growing seasons from sowing to harvest. Phenological phases are as follows: Sowing to Emergence (striped bars), Emergence to BBCH61 (bars without fill pattern), BBCH61 to Physiological Maturity (dotted bars), and Physiological Maturity to Harvest (mesh bars). Dark grey bars indicate the occurrence of the critical period for spring rapeseed according to Kirkegaard et al. (2018). The mean temperature increase due to heat stress treatments is from 0 DAF to 15 DAF (dark grey lines) and between 15 DAF and 30 DAF (light grey lines). The continuous lines illustrate the temperature increase due to heat stress treatments in both genotypes, while dotted lines indicate the temperature increase due to heat stress treatments in only one genotype (either Lumen or Solar CL).

Crop traits were compared using a standard analysis of variance (ANOVA) with a mixed model procedure in Statgraphics Centurion 18. The factor “experiment” was classified as a random term, while the factors “genotype” and “heat stress” were treated as fixed terms. Differences between treatments were considered statistically significant at a probability level of 5%, using the least squares mean differences test. Linear regression analyses were performed using GraphPad Prism 8 to test the fit of the data and the slopes and intercepts of the linear associations. Correlation analyses across yield and its components per experiment were conducted using Statgraphics Centurion 18.

Principal component analysis (PCA) was performed on a data matrix comprising rows that contained combinations of experiments, genotypes, and heat stress treatments (n = 18), and columns that included the following observed variables: grain yield and its components, grain quality (oil and protein concentrations), and climatic variables associated with the treatments recorded during (i) heat stress treatments (0-30 DAF), (ii) grain filling and (iii) the rapeseed grain number critical period. Biplots were constructed using the first two principal components (PC1 and PC2) with JMP 11 from SAS.

The time course of relative individual grain weight and absolute individual grain water content was plotted using GraphPad Prism 8 to explore the impact of the heat stress treatments on individual grain weight. The extra-sum of squares F test and GraphPad Prism 8 were employed for comparisons among the slopes and timings of the end of both the lag and the log phases of the individual grain weight dynamics. The same procedure and software were used to fit the individual grain water content dynamics of grains from the main raceme during grain filling. The normal distribution and homogeneity of residuals were verified across the recorded data (Kutner et al., 2004).

## 3. Results

### 3.1. Environmental conditions and crop phenology across experiments

Air temperature and solar radiation recorded throughout the crop cycles were similar among the experiments, displaying a narrow range (Fig. 2, Table A.1). In the control treatments, the average air temperature between seedling emergence and the start of flowering (BBCH61) ranged from 10.9 to 11.4°C, while during grain filling, it ranged from 14.3°C to 15.3°C (Table A.1). Very similar values were also recorded for average daily solar radiation between emergence and BBCH61 across the experiments (17.2 - 18.9 MJ m^-2^ day^-1^), whereas they were even closer between BBCH61 and physiological maturity (from 24.6 to 24.9 MJ m^2^ day^-1^).

Thermal treatments modified the environmental growing conditions as expected, since portable greenhouse chambers increased the average temperature between 4.1°C and 4.8°C during the 0-15 DAF treatment and from 4.0°C to 5.6°C during the 15-30 DAF (Table 1). Heat load (>30°C) was not recorded in the control treatments, while during the increased temperature 0-15 DAF treatment, heat load ranged from 0 to 48 °Cd and between 16 and 55 °Cd under the 15-30 DAF treatment (Table 1). Average cumulative ISR was slightly lower in the thermal treatments, as the plastic covers in the portable chambers intercepted approximately 10% of the incoming solar radiation (Table 1). Across the experiments, thermal treatments negatively affected the photothermal quotient (PTQ) by decreasing this quotient by 15.5% and 17.9% relative to the controls in the 0-15 DAF and 15-30 DAF treatments, respectively. The PTQ weighted by vapor pressure deficit (PTQ_VPD_) also decreased due to the thermal increase in the 0-15 DAF (31.0%) and 15-30 DAF (31.2%) treatments.

**Table 1.** Mean and maximum temperatures (Tmean and Tmax), cumulative thermal time (TT), temperature differences between heat stress treatments and the control, heat load above 30 °C, incident solar radiation (ISR), cumulative incident solar radiation, photothermal quotient (PTQ) and photothermal quotient corrected for vapour pressure deficit (PTQ_VPD_) during heat stress treatments.

|  |  |  | Tmean | Tmax | Cumulative | Thermal | Heat Load | ISRmean | Cumulative | PTQ | PTQ<sub>VPD</sub> |
| --- | --- | --- | --- | --- | --- | --- | --- | --- | --- | --- | --- |
| Genotype | Moment | Heat Stress | (°C) | (°C) | TT | difference | >30°C | (MJ m<sup>-2</sup> day<sup>-1</sup>) | (MJ m<sup>-2</sup>) | (MJ m<sup>-2</sup> °C<sup>-1</sup>) | (MJ m<sup>-2</sup> °C<sup>-1</sup> Kpa<sup>-1</sup>) |
| Lumen |  |  |  |  |  |  |  |  |  |  |  |
| Exp.1 | 0-15 DAF | C | 13.8 ± 0.5 | 27.2 | 224.6 |  | 0 | 21.4 ± 1.7 | 341.8 | 1.5 ± 0.1 | 3.8 ± 0.5 |
|  |  | H | 18.6 ± 0.5 | 38.2 | 301.2 | 4.8 ± 0.3 | 33 | 19.2 ± 1.6 | 307.6 | 1.0 ± 0.1 | 1.3 ± 0.2 |
|  | 15-30 DAF | C | 14.9 ± 0.4 | 24.8 | 233.1 |  | 0 | 26.9 ± 0.7 | 430.0 | 1.9 ± 0.1 | 3.3 ± 0.3 |
|  |  | H | 18.8 ± 0.4 | 35.4 | 296.3 | 4.0 ± 0.3 | 28 | 24.2 ± 0.6 | 387.0 | 1.3 ± 0.0 | 1.4 ± 0.1 |
| Exp.2 | 0-15 DAF | C | 13.8 ± 0.5 | 27.2 | 225.8 |  | 0 | 22.4 ± 1.7 | 358.7 | 1.6 ± 0.1 | 3.8 ± 0.5 |
|  |  | H | 18.0 ± 0.6 | 39.2 | 292.1 | 4.1 ± 0.3 | 23 | 20.2 ± 1.5 | 322.8 | 1.1 ± 0.1 | 1.5 ± 0.2 |
|  | 15-30 DAF | C | 14.8 ± 0.3 | 24.8 | 230.5 |  | 0 | 26.0 ± 1.2 | 416.2 | 1.8 ± 0.1 | 3.5 ± 0.3 |
|  |  | H | 19.6 ± 0.4 | 35.0 | 308.4 | 4.9 ± 0.4 | 46 | 23.4 ± 1.1 | 374.6 | 1.2 ± 0.1 | 1.2 ± 0.1 |
| Exp.3 | 0-15 DAF | C | 13.2 ± 0.3 | 23.7 | 219.7 |  | 0 | 21.3 ± 2.1 | 362.2 | 1.7 ± 0.2 | 3.6 ± 0.4 |
|  |  | H | 17.6 ± 0.2 | 29.3 | 294.7 | 4.4 ± 0.3 | 0 | 19.2 ± 1.9 | 326.0 | 1.1 ± 0.1 | 1.5 ± 0.1 |
|  | 15-30 DAF | C | 14.3 ± 0.5 | 27.5 | 226.1 |  | 0 | 24.7 ± 1.7 | 395.3 | 1.8 ± 0.1 | 3.2 ± 0.3 |
|  |  | H | 18.2 ± 0.6 | 33.8 | 292.8 | 4.2 ± 0.4 | 21 | 22.2 ± 1.5 | 355.8 | 1.2 ± 0.1 | 1.4 ± 0.1 |
| Solar CL |  |  |  |  |  |  |  |  |  |  |  |
| Exp.1 | 0-15 DAF | C | 14.5 ± 0.6 | 27.2 | 235.1 |  | 0 | 23.0 ± 1.7 | 367.3 | 1.6 ± 0.1 | 3.3 ± 0.5 |
|  |  | H | 18.9 ± 0.6 | 39.7 | 306.7 | 4.5 ± 0.4 | 39 | 20.7 ± 1.5 | 330.6 | 1.1 ± 0.1 | 1.2 ± 0.2 |
|  | 15-30 DAF | C | 14.5 ± 0.3 | 23.8 | 234.8 |  | 0 | 25.4 ± 1.4 | 431.2 | 1.9 ± 0.1 | 3.9 ± 0.3 |
|  |  | H | 19.9 ± 0.4 | 37.9 | 326.6 | 5.6 ± 0.2 | 40 | 22.8 ± 1.2 | 388.1 | 1.2 ± 0.1 | 1.3 ± 0.1 |
| Exp.2 | 0-15 DAF | C | 14.5 ± 0.6 | 27.2 | 235.8 |  | 0 | 24.2 ± 1.5 | 386.6 | 1.6 ± 0.1 | 3.4 ± 0.5 |
|  |  | H | 19.3 ± 0.6 | 37.2 | 312.4 | 4.8 ± 0.3 | 48 | 21.7 ± 1.3 | 347.9 | 1.1 ± 0.1 | 1.3 ± 0.2 |
|  | 15-30 DAF | C | 14.4 ± 0.3 | 22.7 | 205.4 |  | 0 | 24.8 ± 1.5 | 371.7 | 1.8 ± 0.1 | 4.0 ± 0.3 |
|  |  | H | 20.1 ± 0.5 | 36.7 | 291.6 | 5.6 ± 0.4 | 55 | 22.3 ± 1.3 | 334.5 | 1.2 ± 0.1 | 1.3 ± 0.2 |
| Exp.3 | 0-15 DAF | C | 13.4 ± 0.4 | 27.5 | 222.6 |  | 0 | 21.6 ± 2.2 | 366.5 | 1.7 ± 0.2 | 3.5 ± 0.4 |
|  |  | H | 17.7 ± 0.5 | 32.5 | 296.4 | 4.3 ± 0.3 | 3 | 19.4 ± 2.0 | 329.9 | 1.1 ± 0.1 | 1.4 ± 0.2 |
|  | 15-30 DAF | C | 14.2 ± 0.5 | 27.5 | 223.6 |  | 0 | 24.7 ± 1.7 | 395.6 | 1.8 ± 0.1 | 3.3 ± 0.3 |
|  |  | H | 18.9 ± 0.5 | 33.8 | 298.5 | 4.7 ± 0.3 | 16 | 22.3 ± 1.5 | 356.0 | 1.2 ± 0.1 | 1.3 ± 0.1 |

Mean ± standard error

The crop phenology of control plants was similar across genotypes and experiments. For instance, flowering (BBCH61) was reached between 61 and 65 days after sowing, while physiological maturity was recorded from 110 to 130 days after sowing (DAS) across Experiments 1, 2, and 3 (Table A.1). On the other hand, the period between sowing and seedling emergence showed contrasting durations between genotypes in the control plots of Experiments 1 and 2, where seedling emergence was recorded at 9 DAS in Lumen and at 14 DAS in Solar CL. In Experiment 3, both genotypes exhibited similar durations between sowing and seedling emergence, with 17 and 18 DAS in Lumen and Solar CL, respectively. Apparently, no a conclusive reason could be suggested to explain the delays, as the environmental temperature was similar across experiments, as were the sowing depth and soil conditions.

As expected, thermal treatments affected the length of the grain filling period across experiments. Grain filling was shortened by 9, 3, and 6 days in Lumen under the 0-15 DAF treatment in Experiments 1, 2, and 3, respectively, while this period was delayed by 3 and 1 day in Solar CL in Experiments 1 and 2, and shortened by 2 days in Experiment 3 (Fig. A.1). The 15-30 DAF treatment also shortened the grain filling period in Lumen by 11, 3, and 2 days and in Solar CL by 5, 1, and 2 days in Experiments 1, 2, and 3, respectively (Fig. A.1).

### 3.2 Grain yield and above-ground biomass responses to heat stress periods

A wide range of grain yields was recorded in the control treatments, between 4.8 and 7.2 Mg ha^-1^, across genotypes and experiments (Table 2). However, similar grain yields (P > 0.05) were achieved by the controls of Lumen and Solar CL, averaging 5.8 and 6.1 Mg ha^-1^, respectively. The highest yield was recorded in Experiment 3 for both genotypes. Thermal treatments did not show a significant effect (P = 0.061) on grain yield, but an interaction (P = 0.023) between genotype and heat stress treatments was observed across the experiments, where Lumen was less sensitive to higher temperatures than Solar CL (Table 2). The grain yield of both thermal treatments was not associated (P > 0.05) with climatic variables (PTQ, Tmean, Tmax, and cumulative thermal time showed in Table 1) in Lumen, whereas in Solar CL, grain yield was positively associated with PTQ (R^2^ = 0.61; p = 0.023) and negatively with Tmean (R^2^ = 0.61; p = 0.022), Tmax (R^2^ = 0.62; p = 0.021), and cumulative TT (R^2^ = 0.66; p = 0.016). Therefore, Lumen exhibited a grain yield reduction of only 1.2% across heat stress treatments, while this decrease was greater (17.1%) in Solar CL across the experiments and thermal treatments (Table 2). When the periods of heat stress were assessed, the highest impact on grain yield was recorded during the 0-15 DAF treatment. Averaged across the experiments, the grain yield reduction due to the 0-15 DAF treatment reached 16.2%, while during the 15-30 DAF period, this impact was only 2.1%. Between genotypes, Lumen averaged a grain yield reduction of 6.9% in the 0-15 DAF treatment and a negligible increase of 4.0% in the 15-30 DAF treatment. In contrast, Solar CL exhibited a higher reduction in both periods, i.e. 25.5% and 8.2% in the 0-15 and 15-30 DAF treatments, respectively.

**Table 2.**
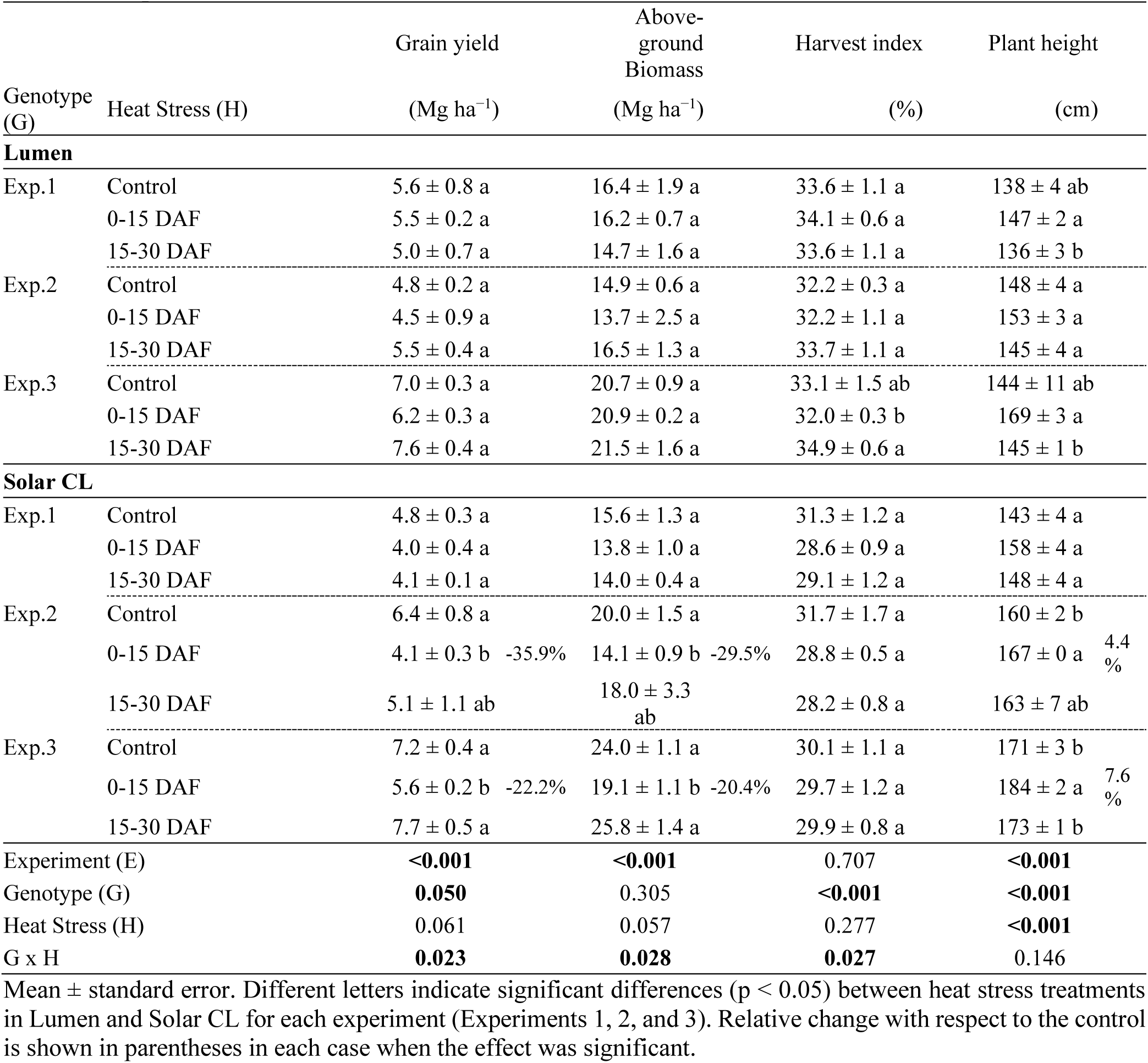
Grain yield, above-ground biomass, harvest index and plant height of the genotypes Lumen and Solar CL in response to heat stress during two periods of grain filling: 0-15 and 15-30 days after flowering (DAF) in Experiments 1, 2 and 3.

|  |  | Grain yield | Above-ground Biomass | Harvest index | Plant height |
| --- | --- | --- | --- | --- | --- |
| Genotype (G) | Heat Stress (H) | (Mg ha<sup>-1</sup>) | (Mg ha<sup>-1</sup>) | (%) | (cm) |
| Lumen |  |  |  |  |  |
| Exp.1 | Control | 5.6 ± 0.8 a | 16.4 ± 1.9 a | 33.6 ± 1.1 a | 138 ± 4 ab |
|  | 0-15 DAF | 5.5 ± 0.2 a | 16.2 ± 0.7 a | 34.1 ± 0.6 a | 147 ± 2 a |
|  | 15-30 DAF | 5.0 ± 0.7 a | 14.7 ± 1.6 a | 33.6 ± 1.1 a | 136 ± 3 b |
| Exp.2 | Control | 4.8 ± 0.2 a | 14.9 ± 0.6 a | 32.2 ± 0.3 a | 148 ± 4 a |
|  | 0-15 DAF | 4.5 ± 0.9 a | 13.7 ± 2.5 a | 32.2 ± 1.1 a | 153 ± 3 a |
|  | 15-30 DAF | 5.5 ± 0.4 a | 16.5 ± 1.3 a | 33.7 ± 1.1 a | 145 ± 4 a |
| Exp.3 | Control | 7.0 ± 0.3 a | 20.7 ± 0.9 a | 33.1 ± 1.5 ab | 144 ± 11 ab |
|  | 0-15 DAF | 6.2 ± 0.3 a | 20.9 ± 0.2 a | 32.0 ± 0.3 b | 169 ± 3 a |
|  | 15-30 DAF | 7.6 ± 0.4 a | 21.5 ± 1.6 a | 34.9 ± 0.6 a | 145 ± 1 b |
| Solar CL |  |  |  |  |  |
| Exp.1 | Control | 4.8 ± 0.3 a | 15.6 ± 1.3 a | 31.3 ± 1.2 a | 143 ± 4 a |
|  | 0-15 DAF | 4.0 ± 0.4 a | 13.8 ± 1.0 a | 28.6 ± 0.9 a | 158 ± 4 a |
|  | 15-30 DAF | 4.1 ± 0.1 a | 14.0 ± 0.4 a | 29.1 ± 1.2 a | 148 ± 4 a |
| Exp.2 | Control | 6.4 ± 0.8 a | 20.0 ± 1.5 a | 31.7 ± 1.7 a | 160 ± 2 b |
|  | 0-15 DAF | 4.1 ± 0.3 b -35.9% | 14.1 ± 0.9 b -29.5% | 28.8 ± 0.5 a | 167 ± 0 a 4.4% |
|  | 15-30 DAF | 5.1 ± 1.1 ab | 18.0 ± 3.3 ab | 28.2 ± 0.8 a | 163 ± 7 ab |
| Exp.3 | Control | 7.2 ± 0.4 a | 24.0 ± 1.1 a | 30.1 ± 1.1 a | 171 ± 3 b |
|  | 0-15 DAF | 5.6 ± 0.2 b -22.2% | 19.1 ± 1.1 b -20.4% | 29.7 ± 1.2 a | 184 ± 2 a 7.6% |
|  | 15-30 DAF | 7.7 ± 0.5 a | 25.8 ± 1.4 a | 29.9 ± 0.8 a | 173 ± 1 b |
| Experiment (E) |  | <0.001 | <0.001 | 0.707 | <0.001 |
| Genotype (G) |  | 0.050 | 0.305 | <0.001 | <0.001 |
| Heat Stress (H) |  | 0.061 | 0.057 | 0.277 | <0.001 |
| G x H |  | 0.023 | 0.028 | 0.027 | 0.146 |

Mean ± standard error. Different letters indicate significant differences ( $p < 0.05$ ) between heat stress treatments in Lumen and Solar CL for each experiment (Experiments 1, 2, and 3). Relative change with respect to the control is shown in parentheses in each case when the effect was significant.

Alike grain yield, above-ground biomass was less affected by heat stress in Lumen than in Solar CL, demonstrating an interaction (P = 0.028) between these sources of variation (Table 2). Across experiments, thermal treatments slightly decreased above-ground biomass in Lumen (0.7%), while in Solar CL this reduction averaged 12.4%. When grain yield was plotted against above-ground biomass across the experiments, a linear and positive association was identified (R^2^ = 0.88; p < 0.001), reinforcing the similar sensitivities between these crop traits (Table 2). Harvest index (HI) was minimally affected by thermal treatments, showing a mean reduction of 0.8 percentage points across genotypes and heat stress treatments; however, there was an interaction (P = 0.027) between genotype and heat stress, where HI of Lumen remained more stable than in Solar CL. Additionally, heat stress increased (P < 0.05) plant height in the 0-15 DAF treatment by 8.2%, with no significant effect (P > 0.05) when temperature was increased during the 15-30 DAF period (Table 2).

### 3.3 Effects of heat stress treatments on grain number and weight

Like grain yield, grain number demonstrated different responses among genotypes and experiments, which is reinforced by the positive association (R^2^ = 0.96; P < 0.001) between these variables when grain yield was plotted against grain number (Fig. 3). Controls of Lumen and Solar CL achieved similar grain number (P > 0.05), i.e. Lumen ranged from 126 to 218 thousand grains per square meter, while Solar between 131 and 212 thousand grains. In Lumen, grain number was affected by heat stress only in Experiment 3, whereas in Solar CL this component was sensitive in Experiments 2 and 3 (Table 3). In both genotypes, the negative impact of increased temperature was recorded under the 0-15 DAF treatment. The highest reduction of grain number was found in Solar CL (−34.8%), while less than half of this reduction (−10.6%) was recorded in Lumen (Table 3). Grain number of Solar CL was positively (R^2^ = 0.61; p = 0.022) associated with photothermal quotient during the critical period for grain number determination of rapeseed reported by Kirkegaard et al. (2018). Across the experiments, grain number was not significantly affected (P > 0.05) by heat stress during the 15-30 DAF period in either Lumen or Solar CL. Regarding TGW, this yield component ranged between 3.14 and 4.29 g across experiments, with a higher TGW was found in Solar CL (3.77 g) than in Lumen (3.61 g), being affected only by genotype. Therefore, grain number was more sensitive to heat stress than thousand grain weight across the experiments (Table 3).

**Figure 3.**
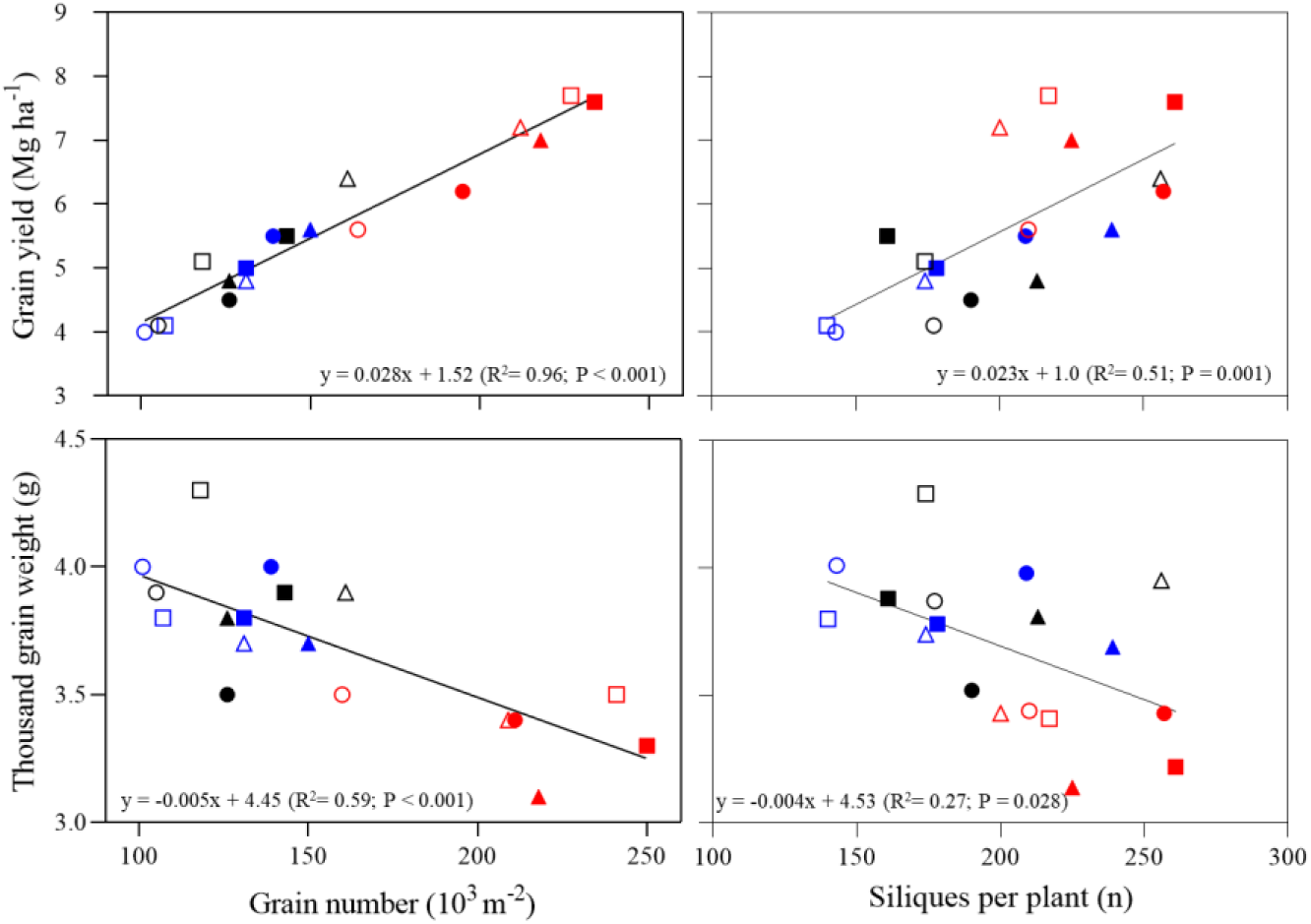
Relationships between grain yield (upper panels) or thousand grain weight (lower panels) and grain number (left panels) and siliques per plant (right panels) of control (triangles), 0-15 DAF (circles) and 15-30 DAF (squares) of genotypes Lumen (closed symbols) and Solar CL (open symbols) in Experiments 1 (blue symbols), 2 (black symbols) and 3 (red symbols).

**Table 3.** Grain number, thousand grain weight and quality traits (oil and protein concentrations) of the genotypes Lumen and Solar CL in response to heat stress during two periods of grain filling: 0-15 and 15-30 days after flowering (DAF) in Experiments 1, 2, and 3.

| Genotype (G) | Heat Stress (H) | Grain number<br>($10 \text{ e}^3 \text{ m}^{-2}$) | TGW<br>(g) | Grain oil concentration<br>(%) | Grain protein concentration<br>(%) |
| --- | --- | --- | --- | --- | --- |
| **Lumen** |  |  |  |  |  |
| Exp.1 | Control | 150 ± 20 a | 3.69 ± 0.07 a | 50.6 ± 0.2 a | 13.8 ± 1.1 a |
|  | 0-15 DAF | 139 ± 6 a | 3.98 ± 0.15 a | 51.1 ± 0.1 a | 14.7 ± 0.7 a |
|  | 15-30 DAF | 131 ± 14 a | 3.78 ± 0.15 a | 50.3 ± 0.8 a | 14.2 ± 0.1 a |
| Exp.2 | Control | 126 ± 7 a | 3.81 ± 0.04 a | 50.5 ± 0.4 a | 15.8 ± 0.1 a |
|  | 0-15 DAF | 126 ± 25 a | 3.52 ± 0.04 b (-7.6%) | 49.9 ± 0.2 a | 15.6 ± 0.4 a |
|  | 15-30 DAF | 143 ± 12 a | 3.88 ± 0.14 a | 49.7 ± 0.5 a | 16.0 ± 0.2 a |
| Exp.3 | Control | 218 ± 5 a | 3.14 ± 0.17 a | 51.0 ± 0.4 ab | 14.9 ± 0.3 a |
|  | 0-15 DAF | 195 ± 1 b (-10.6%) | 3.43 ± 0.05 a | 50.5 ± 0.1 b | 15.2 ± 0.1 a |
|  | 15-30 DAF | 234 ± 13 a | 3.22 ± 0.09 a | 51.7 ± 0.1 a | 15.5 ± 0.2 a |
| **Solar CL** |  |  |  |  |  |
| Exp.1 | Control | 131 ± 14 a | 3.74 ± 0.18 a | 51.0 ± 0.3 a | 15.3 ± 0.4 a |
|  | 0-15 DAF | 101 ± 14 a | 4.01 ± 0.19 a | 50.0 ± 0.6 a | 16.0 ± 0.9 a |
|  | 15-30 DAF | 107 ± 4 a | 3.80 ± 0.03 a | 49.2 ± 0.8 a | 17.0 ± 1.1 a |
| Exp.2 | Control | 161 ± 17 a | 3.95 ± 0.12 ab | 47.8 ± 0.3 a | 17.9 ± 0.2 b |
|  | 0-15 DAF | 105 ± 10 b (-34.8%) | 3.87 ± 0.11 b | 48.6 ± 0.4 a | 18.4 ± 0.4 ab |
|  | 15-30 DAF | 118 ± 22 ab | 4.29 ± 0.12 a | 48.0 ± 0.4 a | 19.5 ± 0.6 a |
| Exp.3 | Control | 212 ± 19 a | 3.43 ± 0.12 a | 50.2 ± 0.3 a | 16.0 ± 0.1 c |
|  | 0-15 DAF | 164 ± 8 b (-22.6%) | 3.44 ± 0.06 a | 48.2 ± 0.7 b (-2.0) | 18.1 ± 0.9 a (2.1) |
|  | 15-30 DAF | 227 ± 17 a | 3.41 ± 0.09 a | 50.2 ± 0.5 a | 16.6 ± 0.2 b (0.6) |
| Experiment (E) |  | **<0.001** | **<0.001** | **<0.001** | **<0.001** |
| Genotype (G) |  | **0.005** | **0.017** | **<0.001** | **<0.001** |
| Heat Stress (H) |  | **0.019** | 0.287 | 0.257 | **0.034** |
| G x H |  | **0.032** | 0.703 | 0.721 | 0.365 |

Mean ± standard error. Different letters indicate significant differences ( $p < 0.05$ ) between heat stress treatments in Lumen and Solar CL for each experiment (Experiments 1, 2, and 3). Relative change with respect to the control is shown in parentheses in each case where the effect was significant. Changes in grain oil and protein with respect to the control are presented as percentage units of increase or reduction, where applicable.

The components of grain number can be divided into siliques per plant and grains per silique. Siliques per plant were only affected by genotype. Averaged across experiments, 27 more siliques per plant were counted in Lumen than in Solar CL (Table 4). When grain yield was plotted against siliques per plant, a positive association (R^2^ = 0.50; P = 0.001) was found (Fig. 3). Additionally, siliques per plant showed a negative trade-off with TGW (R^2^ = 0.27; P = 0.028), meaning that an increase of 100 siliques per plant reduced TGW by 0.4 g.

**Table 4.** Siliques per plant, weight per silique, grains per silique, and branch number of the genotypes Lumen and Solar CL in response to heat stress during two periods of grain filling: 0-15 and 15-30 days after flowering (DAF) in Experiments 1, 2 and 3.

| Genotype (G) | Heat Stress (H) | Siliques per plant (n) | Weight per silique (mg silique<sup>-1</sup>) | Grains per silique (n silique<sup>-1</sup>) | Branch number (n per plant) |
| --- | --- | --- | --- | --- | --- |
| **Lumen** |  |  |  |  |  |
| Exp.1 | Control | 239 ± 6 a | 41 ± 5 a | 11.0 ± 1.3 a | 7.6 ± 0.4 a |
|  | 0-15 DAF | 209 ± 33 a | 49 ± 8 a | 12.1 ± 1.8 a | 6.6 ± 0.6 a |
|  | 15-30 DAF | 178 ± 41 a | 57 ± 21 a | 14.8 ± 4.8 a | 6.8 ± 0.3 a |
| Exp.2 | Control | 213 ± 11 a | 40 ± 3 a | 10.4 ± 1.0 a | 7.0 ± 0.6 a |
|  | 0-15 DAF | 190 ± 36 a | 44 ± 12 a | 12.6 ± 3.5 a | 6.2 ± 0.4 a |
|  | 15-30 DAF | 161 ± 26 a | 64 ± 11 a | 16.6 ± 3.3 a | 6.4 ± 0.4 a |
| Exp.3 | Control | 225 ± 13 a | 53 ± 2 a | 17.0 ± 0.6 a | 6.3 ± 0.2 a |
|  | 0-15 DAF | 257 ± 18 a | 44 ± 3 b (-17.0%) | 12.9 ± 0.7 b (-24.1%) | 6.0 ± 0.1 a |
|  | 15-30 DAF | 261 ± 24 a | 51 ± 1 ab | 15.8 ± 0.6 a | 6.2 ± 0.4 a |
| **Solar CL** |  |  |  |  |  |
| Exp.1 | Control | 174 ± 18 a | 49 ± 5 a | 13.4 ± 1.8 a | 6.4 ± 0.1 a |
|  | 0-15 DAF | 143 ± 14 a | 50 ± 9 a | 12.8 ± 2.7 a | 5.9 ± 0.2 a |
|  | 15-30 DAF | 140 ± 35 a | 57 ± 12 a | 14.9 ± 3.1 a | 5.9 ± 0.3 a |
| Exp.2 | Control | 256 ± 46 a | 50 ± 18 a | 12.5 ± 4.1 a | 6.3 ± 0.4 a |
|  | 0-15 DAF | 177 ± 22 a | 42 ± 9 a | 11.0 ± 2.6 a | 5.9 ± 0.3 a |
|  | 15-30 DAF | 174 ± 10 a | 52 ± 12 a | 12.1 ± 2.4 a | 6.3 ± 0.2 a |
| Exp.3 | Control | 200 ± 7 a | 63 ± 4 a | 18.6 ± 2.0 a | 6.3 ± 0.1 a |
|  | 0-15 DAF | 210 ± 32 a | 49 ± 6 a | 14.3 ± 2.0 a | 5.7 ± 0.2 b (-9.5%) |
|  | 15-30 DAF | 217 ± 16 a | 62 ± 1 a | 18.3 ± 0.4 a | 6.4 ± 0.1 a |
| Experiment (E) |  | **0.013** | 0.424 | **0.017** | 0.219 |
| Genotype (G) |  | **0.041** | 0.537 | 0.915 | **0.013** |
| Heat Stress (H) |  | 0.172 | 0.204 | 0.248 | **0.011** |
| G x H |  | 0.680 | 0.569 | 0.622 | 0.645 |

Mean ± standard error. Different letters indicate significant differences ( $p < 0.05$ ) between heat stress treatments in Lumen and Solar CL for each experiment (Experiments 1, 2, and 3). Relative change with respect to the control is shown in parentheses in each case when the effect was significant.

Interestingly, individual silique weight and grains per silique were not affected by genotype or thermal treatments, while the number of branches per plant was influenced by both sources of variation (Table 4). Branch number per plant in Lumen and Solar CL averaged 6.6 and 6.1, respectively. Across experiments, thermal treatment during the 0-15 DAF period reduced (P < 0.05) branch number by 10% and 7.9% in Lumen and Solar CL, respectively, while no impact (P > 0.050) of heat stress was found during the 15-30 DAF heat treatment. Regarding TGW, when the time course of individual grains was recorded, the weight and water content of grains from the main raceme were mainly affected by genotype, but little sensitivity of individual grain dynamics was observed due to heat treatments in the experiments (Fig. A.1).

### 3.4 Sensitivity of quality traits (grain oil and protein concentration) to heat stress

Oil concentration of grains ranged from 47.8% to 51.7%, and protein concentration varied between 13.8% and 19.5%. As expected, grain oil concentration was relatively conservative despite the different grain yields recorded in both genotypes across the experiments (Table 3 and Fig A.3). In the control treatments, Lumen and Solar CL reached similar oil concentrations in grains (50.7% and 49.7%, respectively). This trait was only affected by heating in Experiment 3 (Table 3). On the other hand, grain protein concentration was sensitive to both genotype and thermal treatments, showing an increase of 2.1 percentage points in Solar CL compared to Lumen when averaged across heat treatments. Regarding grain protein concentration, both thermal treatments had a positive effect (P < 0.05) on this trait, increasing it by 2.2 to 6.7 percentage points in the 0-15 DAF treatment and from 2.7 to 7.9 percentage points in the 15-30 DAF treatment relative to the control, averaged across genotypes and experiments. A trade-off between grain protein and oil concentrations was observed across the recorded data (Fig. 4).

**Figure 4.**
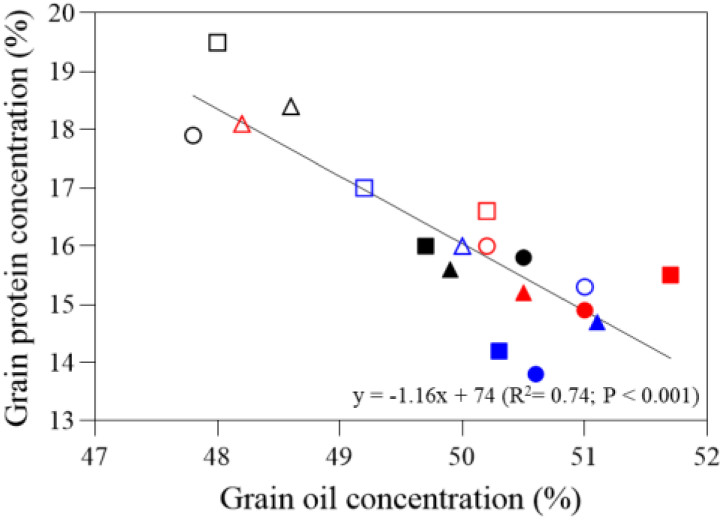
Relationship between grain oil and protein concentrations of control (triangles), 0-15 DAF (circles), and 15-30 DAF (squares) treatments of Lumen (closed symbols) and Solar CL (open symbols) in Experiments 1 (blue symbols), 2 (black symbols) and 3 (red symbols).

### 3.5 Analysis of grain yield, quality traits and environmental conditions across the experiments

An integrated PCA analysis was performed, including crop responses and environmental variables recorded during (i) heat stress treatments (0-30 DAF), (ii) grain filling and (iii) the rapeseed grain number critical period (Fig. 5). Components PC1 and PC2 explained 70.5% of the total variation. Component PC1 primarily discriminated trade-offs, i.e., (i) grain number and thousand grain weight or (ii) grain oil and protein concentration. In contrast, component PC2 discriminated between Experiments 1 and 2 against Experiment 3.

**Figure 5.**
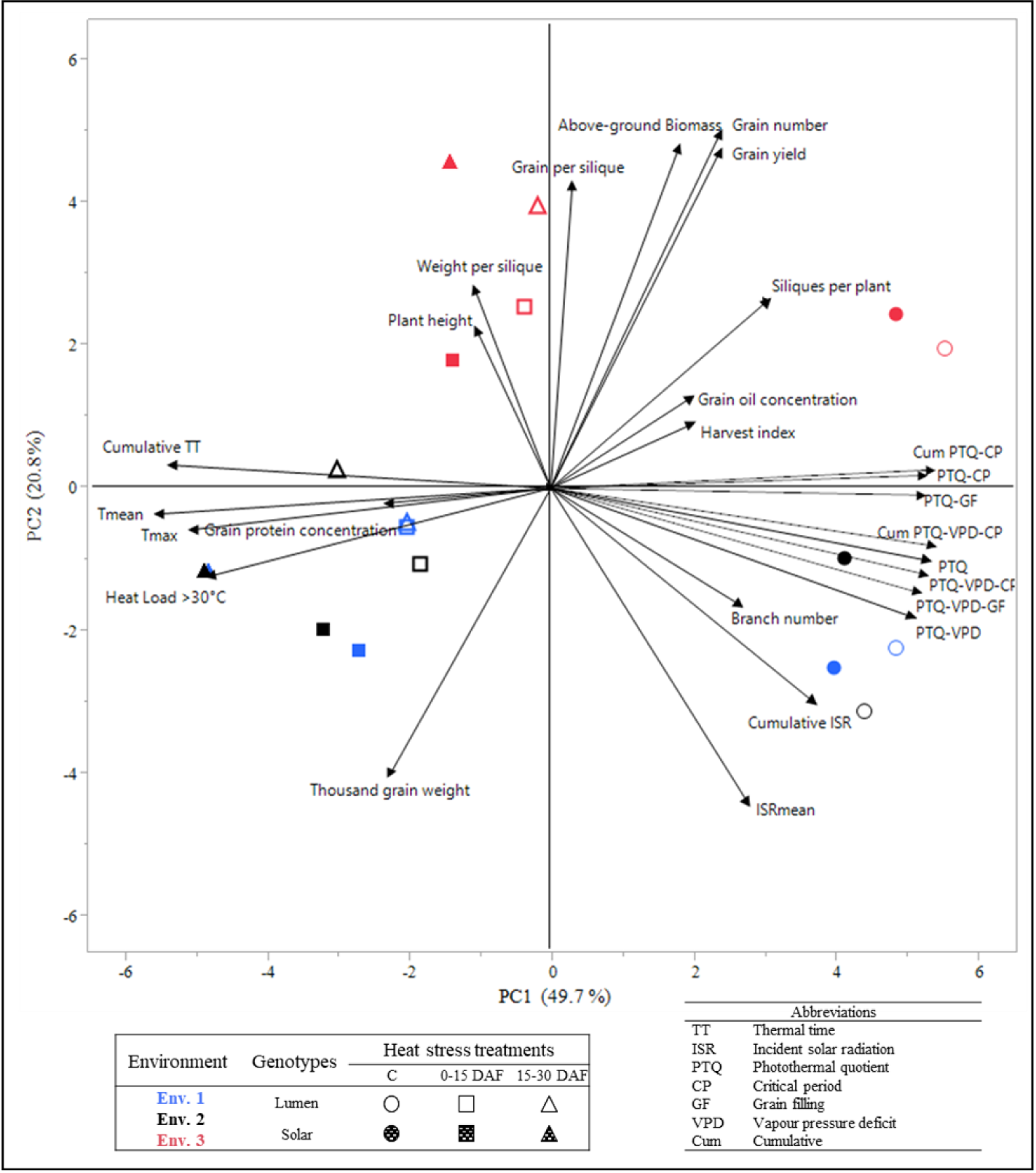
Principal component analysis (PCA) of grain yield components and weather data of control (circles), 0-15 DAF (squares), and 15-30 DAF (triangles) treatments of Lumen (open symbols) and Solar CL (closed symbols) in Experiments 1 (blue symbols), 2 (black symbols) and 3 (red symbols). The two axes explained 70.5% of the variation.

In agreement with the analysis reported above, grain yield was positively associated with above-ground biomass and grain number, while negatively correlated with TGW. Environmental conditions favouring grain yield in Experiments 1 and 3 were associated with photothermal quotients (PTQ) during grain filling (PTQ-GF) or throughout the critical period (PTQ-CP). Conversely, temperatures (Tmean, Tmax, and cumulative TT) and heat load were negatively associated with the photothermal quotients. The genotypes showed different associations with the environmental variables; Lumen was not associated with climatic variables, whereas Solar CL was (Fig. A.2). Additionally, grain number and silique number per plant were negatively associated with Tmean and heat load while being positively correlated with TGW. Siliques per plant were also positively correlated with PTQ-GF and PTQ-CP (R^2^ = 0.22, P = 0.050 and R^2^ = 0.23, P = 0.042, respectively). ISR was negatively associated with grain number and grains per silique but positively associated with TGW and branch number. Quality traits showed no correlation with climate variables.

## 4. Discussion

### 4.1 Sensitivity of grain yield to heat stress after flowering

The objective of the present study was to assess the sensitivity of rapeseed to the temperature increase at two windows of time during grain filling, i.e., 0-15 and 15-30 DAF. In a previous study, different sensitivity of rapeseed was found between these phases of the grain filling period under the reduction of the source-sink ratio (Verdejo and Calderini, 2020). Additionally, partial or complete resilience of grain yield to the source shortage during grain filling have been reported in this crop due to the plasticity of grain weight (Kirkegaard et al., 2018; Labra et al., 2017; Verdejo and Calderini, 2020). Therefore, we aimed to evaluate the behaviour of rapeseed under another abiotic constraint, which is forecasted in the scenarios of climate change, i.e. heat stress.

In our experiment, climate conditions of control treatments agree with the average monthly temperature and radiation of the historical records for Valdivia and southern Chile (Mera et al., 2015; Verdejo and Calderini, 2020). Grain yield of control treatments in Lumen and Solar CL genotypes confirmed the high-yielding conditions already reported for temperate crops in southern Chile through different experiments (Bustos et al., 2013; del Pozo et al., 2022; Rivelli et al., 2024). In addition, the average temperature across the whole assessed period of the control treatments ranged from 13.2 to 14.9°C, while they were between 16.0 and 16.9°C in the 0-15 DAF and from 15.7 to17.3°C in the 15-30 DAF treatments (Table 1). Therefore, the magnitude of the heat increment in our experiments averaged 4.5 °C in the 0-15 DAF and 4.8°C in the 15-30 DAF periods over the control. The mean maximum temperature recorded across our experiments (up to 20.1 °C) was lower than most of the previous studies reaching between 23.7 and 34 °C (Elferjani and Soolanayakanahally, 2018; Gan et al., 2004; Morrison, 1993; Pokharel et al., 2021). Thus, although a temperature increment of 4°C proposed in our study is realistic four Southern Chile, the background and maximum temperatures of our experiments were different from other studies carried out on rapeseed in different environmental conditions (Lee et al., 2021).In addition, control treatments of the above mentioned studies recorded higher mean temperatures, i.e. from 19.3 to 20.3 °C (Elferjani and Soolanayakanahally, 2018; Gan et al., 2004; Morrison, 1993; Pokharel et al., 2021). The sensitivity of rapeseed to increased temperature evaluated in most of the previous studies was carry out under controlled conditions (Elferjani and Soolanayakanahally, 2018; Gan et al., 2004; Morrison, 1993; Pokharel et al., 2021; Rao et al., 1992; Secchi et al., 2023), standing out the importance of the present study under field conditions.

High temperature early after flowering showed a negative impact on grain yield of rapeseed in previous assessments performed using historical weather and crop yield data from 1967 to 2001, along different crop districts in Saskatchewan, Canada (Kutcher et al., 2010). Additionally, in a meta-analysis by Secchi et al. (2023) the authors reported that yield reduction due to heat stress depends on the phenological period when the stress occurs and the magnitude of the temperature increase. These authors showed that heat stress before the end of flowering caused grain yield reductions ranging between 20 and 39%, while after the end of flowering grain yield decreased ∼21%. In our experiments, slightly effects were found due to thermal increase on grain yield (Table 2). But both hybrids evaluated in our experiments showed different response to heat stress, i.e. grain yield was less affected by heating in Lumen (P > 0.05) than in Solar CL (P < 0.05). This differential response of genotypes might be due to the later released hybrid Lumen developed under new breeding approach attempting to breed climate-smart genotypes (Varshney et al., 2021). In this way, Lumen and Solar CL were release in Denmark in 2015 and 2011, respectively (Deutsche Saatveredelung AG, personal communication). Regarding crop traits, the higher grain yield resilience showed by Lumen could be due to a compensation by later-formed siliques during grain filling (Table 4 and Fig. 4) respect to Solar CL (Aksouh-Harradj et al., 2006; Labra et al., 2017; Massuia de Almeida et al., 2021; Zheng et al., 2022). Additionally, Canales et al. (2023) reported a genotype dependent variation in the regulation of photosynthetic genes of Lumen and Solar CL under heat stress, which may confer thermotolerance in Lumen by sustained carbon assimilation.

### 4.2 Grain number and weight under heat stress after flowering

As expected, grain number was the main determinant of grain yield across genotypes, thermal treatments and experiments (R^2^= 0.94; P< 0.001), while grain weight showed a negative association (R^2^= 0.42; P= 0.004) (Fig. 3) due to the often found trade-off between both yield components in rapeseed (Rivelli et al., 2023; Verdejo and Calderini, 2020) and other crops (e.g. Achary and Reddy, 2021; Acreche and Slafer, 2006; Brunner et al., 2024; Calderini et al., 2021). Interestingly, grain number of Lumen was not affected in most of the experiments and heat stress periods showing the same degree of resilience than grain yield. On the other hand, grain number was statistically affected by the increased temperature in Solar CL in Experiments 2 and 3, though contrasting sensitivity of this trait was found between heat stress periods (Table 3). For instance, raised temperature had a greater negative impact on grain number of Solar CL in the 0-15 DAF treatment than during the 15-30 DAF, i.e., grain number reductions were 28.7% and 9.9%, for these periods across Experiments 2 and 3, respectively. It is important to highlight that the heat stress periods assessed in the present study (0-15 and 15-30 DAF) overlap with the rapeseed critical period (100-500 °Cd after flowering) for grain number determination found by Kirkegaard et al. (2018). This overlap accounted for 56% in the 0-15 DAF and 51% during the 15-30 DAF period, which were similar to the overlapping of the source-sink reduction carried out in our previous study (Verdejo and Calderini, 2020). This allows us to compare both abiotic stresses and windows time.

The results reported above are different from the previous study evaluating the response of grain number of the same rapeseed hybrids to the source-sink reduction carried out in Valdivia, where similar sensitivity of grain number was reported during both 0-15 DAF and 15-30 DAF periods (Verdejo and Calderini, 2020). Specifically, grain number reductions in Solar CL were 46.4 and 34.7.% in the 0-15 and 15-30 DAF, respectively, under source-sink ratio reduction, however, in that study TGW increased 35.6% in the 0-15 DAF treatment, compensating therefore, the GN decrease, allowing grain yield resilience (Verdejo and Calderini, 2020). The present study shows different behaviour under increased temperature, at least in the high yielding environment of southern Chile. Additionally, the sensitivity of rapeseed to both abiotic stresses during the 0-15 DAF period was consistent (Verdejo and Calderini, 2020), but depended on the genotype. In agreement with our results, Rivelli et al. (2024) assessing Solar CL in Valdivia, also found a clear tolerance of grain number to heat stress when the temperature was increased by 5 °C from 10 to 20 DAF in this hybrid.

Contrary to previous heat stress studies (Aksouh et al., 2001; Gan et al., 2004; Pokharel et al., 2021; Triboi-Blondel and Renard, 1999), both genotypes assessed in our experiments (Lumen and Solar CL) showed very little impact of heat stress on grain weight during both 0-15 DAF and 15-30 DAF periods (Table 3). The grain weight tolerance found in our study is of high interest due to the contrasting results reported in the literature. For example, heat stress studies have shown grain weight reduction of 19% (Triboi-Blondel and Renard, 1999) when temperature was increased 7.3 °C during the grain filling period. Specifically, Triboi-Blondel and Renard (1999) performed heat stress treatments from late flowering to maturity assessing a winter type rapeseed genotype. Additionally, heat stress studies commonly reported both grain number and grain weight reductions. For instance, Gan et al. (2004) showed similar reduction of 22% of both grain number and thousand grain weight when the authors contrasted a heat stress regime of 35/18 °C with a control at 20/18°C (day/night temperature) resulting in a 38% grain yield reduction.

Differences among studies showing or not negative impact of higher temperature during grain filling on grain yield components could be explained by a lower intensity of the heat stress compared with other studies as commented above (Elferjani and Soolanayakanahally, 2018; Gan et al., 2004; Morrison, 1993; Pokharel et al., 2021). Therefore, the temperature of 34 °C pointed out as the threshold for deleterious grain yield effect (Hatfield et al., 2011) was not reached in our experiments (Table 1 and A.1). Additionally, the background temperature in southern Chile (Mera et al., 2015) might allow a major tolerance of grain yield to increased temperature as it was pointed out for other crops (García et al., 2015). For example, Sadras and Moran (2013) found that the hotter the environment the higher the heat stress impact on grapevine.

The molecular mechanisms involved in the rapeseed response to heat stress of our study were addressed by Canales et al. (2023). The authors discovered novel gene modules regulating the complex physiological responses of grain number and grain weight to heat stress in rapeseed during the critical reproductive stages of flowering and early seed set. They found that genes involved in grain weight development were associated with biological functions related to intracellular transport, mRNA metabolism, cell signalling, and chromatin remodelling. In addition, the transcription factors INO (INNER NO OUTER), ABI4 (ABA INSENSITIVE 4) and ARF2 (AUXIN RESPONSE FACTOR 2) seem to be key for grain number set and grain development at the early grain filling period. Therefore, the complement between physiological and molecular approaches could provide clues for a better understanding of the response of rapeseed to heat, likely explaining the resilience of this crop in southern Chile. For example, combining low background and maximum temperatures, low heat load and genes of the transcription factors INO, ABI4 and chromatin remodelling.

### 4.3 Sensitivity of rapeseed grain quality traits to heat stress

Heat stress treatments showed no effect on grain oil concentration and a positive effect was found on grain protein concentration in the present study (Table 3), which contrast with some previous heat stress studies on rapeseed. Heat stress often has negative effects on oil grain concentration and positive effects on protein grain concentration in rapeseed (Faraji, 2012; Zhu et al., 2012). For instance, (Pokharel et al., 2021); Triboi-Blondel and Renard (1999) found a decrease of 6 percentage points in the grain oil concentration and an increase of 7 percentage points in grain protein concentration under controlled conditions. Similarly, Pokharel et al. (2021) reported a decrease in grain oil concentrations of 3.2 percentage points and a grain protein concentration increase of 3.7 percentage points in field conditions when temperature was increased up to 34°C seven days after 50% of flowering. Although a trade-off between grain protein and oil concentration was found in our study (Fig. 4) as previously (Kirkegaard et al., 2018; Labra et al., 2017; Rondanini et al., 2014) grain quality traits showed much higher stability in our experiments than grain yield, grain number and thousand grain weight (Fig A.3). The stability of grain oil concentration under heat stress was in accordance with previous studies in southern Chile evaluating source-sink reductions and heat stress treatments in oil crops such as sunflower and rapeseed where grain oil concentration was almost not affected by the treatments (Castillo et al., 2017; Labra et al., 2017; Rivelli et al., 2024). In addition, Poudel et al. (2024) found in soybean that grain oil concentration is more stable than protein grain concentration when a pod removal treatment was assessed, in agreement to our study despite high differences between both crop species.

## 5. Conclusions

Results reported in this study show slightly effects due to thermal increase on grain yield across experiments in the high yield environment of southern Chile. The main impact of heat stress on grain yield after flowering was during the first half of grain filling, especially through grain number reduction, but the sensitivity of this trait depended on the genotype as the Lumen hybrid was clearly less affected than Solar CL. The impact of temperature increase during the 0-15 DAF agrees with previous studies performed with the same genotype under source-sink reduction. On the other hand, heat stress scarcely affected grain weight in southern Chile. The lack of sensitivity of grain weight under both timings of the thermal stress (0-15 and 15-30 DAF) is of high importance due to the showed tolerance of this trait recorded in southern Chile respect to other heat stress studies assessing rapeseed response. Finally, grain oil was almost not affected by the temperature increase after the start of flowering, being this key trait highly conservative across genotypes and stress periods, while grain protein concentration was more sensitive, increasing under heat stress.

## 6. Author contribution

José F. Verdejo: Conceptualization, Formal analysis, Investigation, Writing - original draft, Visualization. Daniel F. Calderini.: Conceptualization, Methodology, Resources, Supervision, Writing - review & editing. All authors read and approved the final version of the manuscript.

## 7. Acknowledgements

We thank Olaff Sass (NPZ-Lembke, Germany) and Erik von Baer (Semillas Baer, Chile) for kindly providing the seeds of the genotypes. We really appreciate technical support from the staff of the Austral Farming Experimental Station (EEAA) of the Universidad Austral de Chile. The present study was supported by Project FONDECYT 1170913 (Chilean Technical and Scientific Research Council, CONICYT/ANID) competitive grant. José Verdejo held a postgraduate scholarship from CONICYT (ANID) 2017-21171384.

---

**Table A.1.**
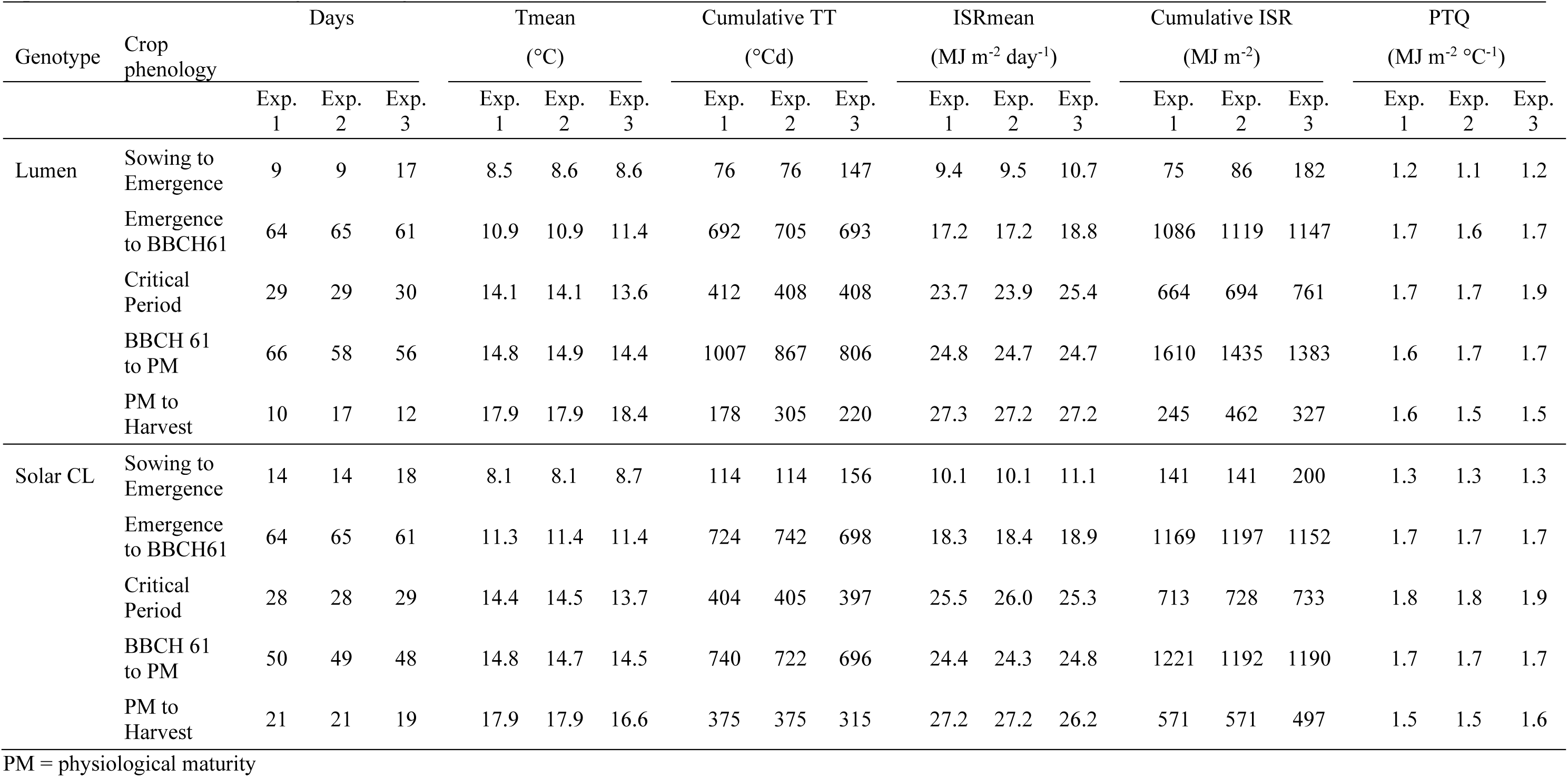
Days, mean temperatures (Tmean), cumulative thermal time (TT), incident solar radiation (ISR), cumulative incident solar radiation and photothermal quotient (PTQ) throughout the crop phenology of Lumen and Solar CL in control treatments. The critical period for spring rapeseed is shown according to Kirkegaard et al. (2018).

**Figure A.1.**
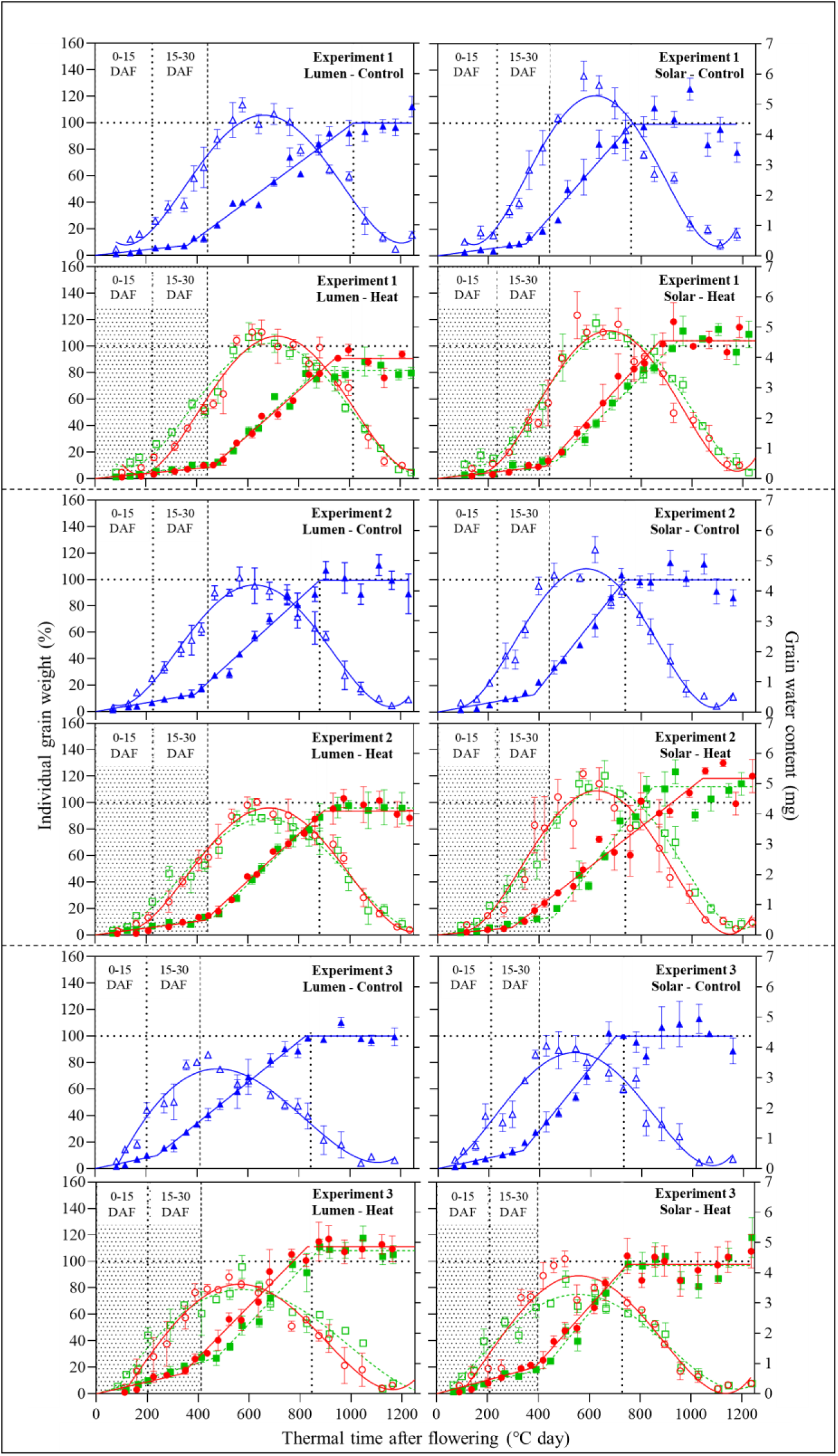
Time course of individual grain weight and grain water content of seeds from the main raceme of the genotypes Lumen (left panels) and Solar CL (right panels) under control (open and closed triangles), 0-15 DAF (open and closed circles) and 15-30 DAF (open and closed squares) heat stress treatments. Individual grain weight is expressed relative to the control. Bars indicate the standard error of the means. Shaded areas represent the timing of both the 0-15 and 15-30 DAF heat stress treatments, while dotted lines mark the moment of physiological maturity for the control treatments.

**Figure A.2.**
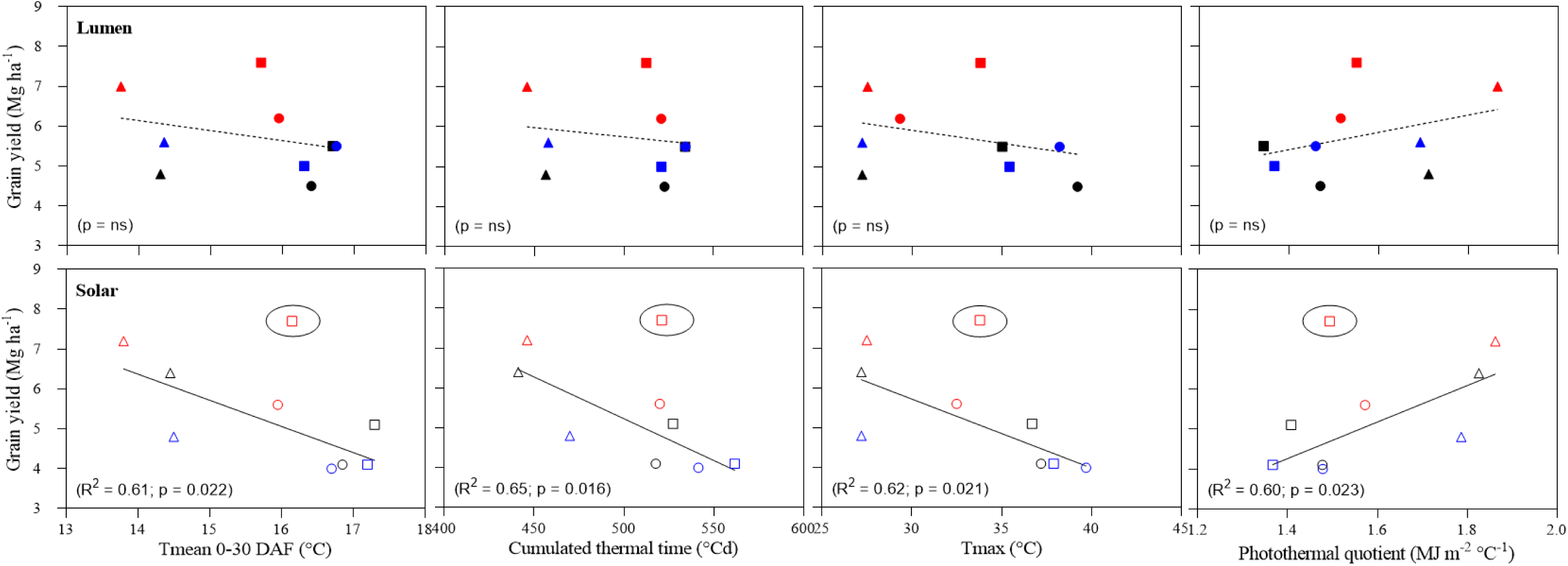
Relationships between grain yield and temperature variables during the imposed heat stress treatments (0-30 days after flowering) for Lumen (upper panels and closed symbols) and Solar CL (lower panels and open symbols) for control (triangles), 0-15 DAF (circles) and 15-30 DAF (squares) in Experiments 1 (blue symbols), 2 (black symbols) and 3 (red symbols). The circled red square in Solar CL was excluded from the regression analysis as an outlier.

**Fig A.3.**
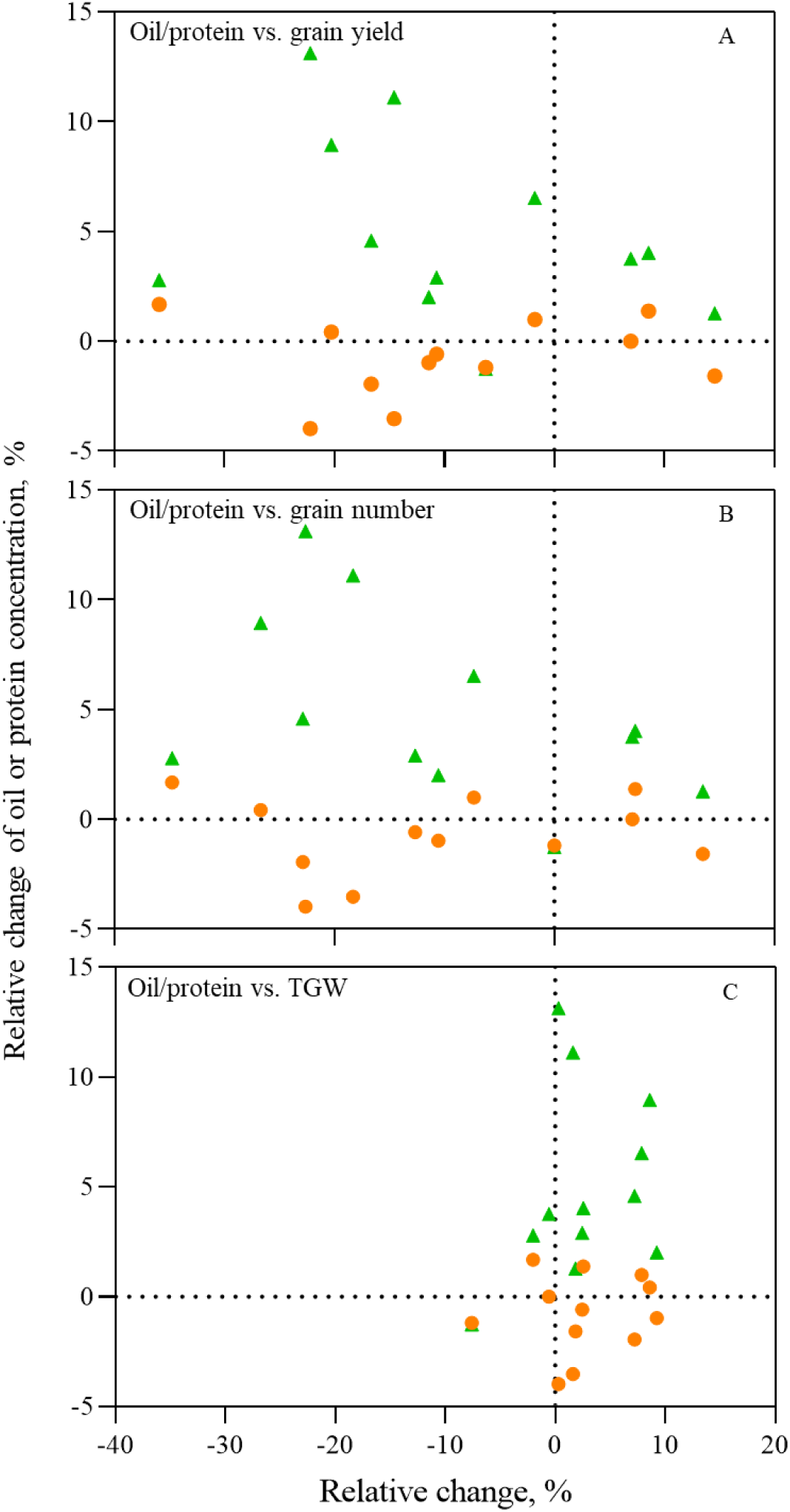
Association between the relative change of oil (orange circles) or protein (green triangles) concentrations in grains and the relative change of grain yield (A), grain number (GN) (B), and thousand grain weight (TGW) (C) across the experiments.

## Notes

### Competing Interest Statement

The authors have declared no competing interest.

## 8. References

Achary, V.M.M. and Reddy, M.K., 2021. CRISPR-Cas9 mediated mutation in GRAIN WIDTH and WEIGHT2 (GW2) locus improves aleurone layer and grain nutritional quality in rice. Sci Rep-Uk, 11(1): 21941. 10.1038/s41598-021-00828-z

Acreche, M.M. and Slafer, G.A., 2006. Grain weight response to increases in number of grains in wheat in a Mediterranean area. Field Crop Res, 98(1): 52–59. 10.1016/j.fcr.2005.12.005

Aksouh-Harradj, N.M., Campbell, L.C. and Mailer, R.J., 2006. Canola response to high and moderately high temperature stresses during seed maturation. Can J Plant Sci, 86(4): 967–980. 10.4141/p05-130

Aksouh, N.M., Jacobs, B.C., Stoddard, F.L. and Mailer, R.J., 2001. Response of canola to different heat stresses. Aust J Agr Res, 52(8): 817–824. 10.1071/AR00120

Araya-Osses, D., Casanueva, A., Román-Figueroa, C., Uribe, J.M. and Paneque, M., 2020. Climate change projections of temperature and precipitation in Chile based on statistical downscaling. Climate Dynamics, 54(9): 4309–4330. 10.1007/s00382-020-05231-4

Asseng, S. et al., 2015. Rising temperatures reduce global wheat production. Nature Climate Change, 5(2): 143–147. 10.1038/nclimate2470

Awasthi, R. et al., 2014. Individual and combined effects of transient drought and heat stress on carbon assimilation and seed filling in chickpea. Funct Plant Biol, 41(11): 1148–1167. 10.1071/FP13340

Bonada, M. and Sadras, V.O., 2015. Review: critical appraisal of methods to investigate the effect of temperature on grapevine berry composition. Aust J Grape Wine R, 21(1): 1–17. 10.1111/ajgw.12102

Brunner, S., Weichert, H., Meissle, M., Romeis, J. and Weber, H., 2024. Field trials reveal trade-offs between grain size and grain number in wheat ectopically expressing a barley sucrose transporter. Field Crop Res, 316: 109506. 10.1016/j.fcr.2024.109506

Bustos, D.V., Hasan, A.K., Reynolds, M.P. and Calderini, D.F., 2013. Combining high grain number and weight through a DH-population to improve grain yield potential of wheat in high-yielding environments. Field Crop Res, 145: 106–115. 10.1016/j.fcr.2013.01.015

Calderini, D.F. et al., 2021. Overcoming the trade-off between grain weight and number in wheat by the ectopic expression of expansin in developing seeds leads to increased yield potential. New Phytol, 230(2): 629–640. 10.1111/nph.17048

Campbell, G.S., 1981. Fundamentals of Radiation and Temperature Relations. In: O.L. Lange, P.S. Nobel, C.B. Osmond and H. Ziegler (Editors), Physiological Plant Ecology I: Responses to the Physical Environment. Springer Berlin Heidelberg, Berlin, Heidelberg, pp. 11–40.

Canales, J., Verdejo, J.F. and Calderini, D.F., 2023. Transcriptome and physiological analysis of rapeseed tolerance to post-flowering temperature increase. Int J Mol Sci, 24(21). 10.3390/ijms242115593

Castillo, F.M., Vásquez, S.C. and Calderini, D.F., 2017. Does the pre-flowering period determine the potential grain weight of sunflower? Field Crop Res, 212: 23–33. 10.1016/j.fcr.2017.06.029

Cheng, W., Sakai, H., Yagi, K. and Hasegawa, T., 2009. Interactions of elevated [CO2] and night temperature on rice growth and yield. Agr Forest Meteorol, 149(1): 51–58. 10.1016/j.agrformet.2008.07.006

Cossani, C.M. and Sadras, V.O., 2021. Nitrogen and water supply modulate the effect of elevated temperature on wheat yield. Eur J Agron, 124: 126227. 10.1016/j.eja.2020.126227

del Pozo, A. et al., 2022. Genetic yield gains and changes in morphophysiological-related traits of winter wheat in southern chilean high-yielding environments. Front Plant Sci, 12. 10.3389/fpls.2021.732988

Egli, D.B., 2017. Seed Biology and the Yield of Grain Crops. CAB International, Wallingford, Oxfordshire; Boston, MA, 234 pp.

Egli, D.B., TeKrony, D.M., Heitholt, J.J. and Rupe, J., 2005. Air temperature during seed filling and soybean seed germination and vigor. Crop Sci, 45(4): 1329–1335. 10.2135/cropsci2004.0029

Elferjani, R. and Soolanayakanahally, R., 2018. Canola responses to drought, heat, and combined stress: shared and specific effects on carbon assimilation, seed yield, and oil composition. Front Plant Sci, 9. 10.3389/fpls.2018.01224

Faraji, A., 2012. Oil concentration in canola (Brassica napus L.) as a function of environmental conditions during seed filling period. Int J Plant Prod, 6(3): 267–277. 10.22069/ijpp.2012.764

Gan, Y. et al., 2004. Canola and mustard response to short periods of temperature and water stress at different developmental stages. Can J Plant Sci, 84(3): 697–704. 10.4141/p03-109

García, G.A., Dreccer, M.F., Miralles, D.J. and Serrago, R.A., 2015. High night temperatures during grain number determination reduce wheat and barley grain yield: a field study. Global Change Biology, 21(11): 4153–4164. 10.1111/gcb.13009

Hatfield, J.L. et al., 2011. Climate impacts on agriculture: implications for crop production. Agron J, 103(2): 351–370. 10.2134/agronj2010.0303

Hernandez, K. and Madeira, C., 2022. The impact of climate change on economic output across industries in Chile. Plos One, 17(4): e0266811. 10.1371/journal.pone.0266811

Kim, H.-R. and You, Y.-H., 2010. Effects of elevated CO2 concentration and increased temperature on leaf related-physiological responses of Phytolacca insularis (native species) and Phytolacca americana (invasive species). Journal of Ecology and Environment, 33(3): 195–204. 10.5141/jefb.2010.33.3.195

Kirk, P.L., 1950. Kjeldahl method for total nitrogen. Analytical Chemistry, 22(2): 354–358. 10.1021/ac60038a038

Kirkegaard, J.A., Lilley, J.M., Brill, R.D., Ware, A.H. and Walela, C.K., 2018. The critical period for yield and quality determination in canola (Brassica napus L.). Field Crop Res, 222: 180–188. 10.1016/j.fcr.2018.03.018

Kirkegaard, J.A. et al., 2012. Physiological response of spring canola (Brassica napus) to defoliation in diverse environments. Field Crop Res, 125: 61–68. 10.1016/j.fcr.2011.08.013

Konuskan, D.B., Arslan, M. and Oksuz, A., 2019. Physicochemical properties of cold pressed sunflower, peanut, rapeseed, mustard and olive oils grown in the Eastern Mediterranean region. Saudi Journal of Biological Sciences, 26(2): 340–344. 10.1016/j.sjbs.2018.04.005

Kutcher, H.R., Warland, J.S. and Brandt, S.A., 2010. Temperature and precipitation effects on canola yields in Saskatchewan, Canada. Agr Forest Meteorol, 150(2): 161–165. 10.1016/j.agrformet.2009.09.011

Kutner, M., Nachtsheim, C. and Neter, J., 2004. Applied Linear Regression Models. McGraw-Hill Education, Boston, 701 pp.

Labra, M.H., Struik, P.C., Evers, J.B. and Calderini, D.F., 2017. Plasticity of seed weight compensates reductions in seed number of oilseed rape in response to shading at flowering. Eur J Agron, 84(Supplement C): 113–124. 10.1016/j.eja.2016.12.011

Lambers, H. and Oliveira, R.S., 2019. Plant Physiological Ecology. Springer International Publishing, XXVII, 736 pp. 10.1007/978-3-030-29639-1

Lee, J.-Y. et al., 2021. Future Global Climate: Scenario-Based Projections and Near-Term Information. In: V. Masson-Delmotte et al. (Editors), Climate Change 2021: The Physical Science Basis. Contribution of Working Group I to the Sixth Assessment Report of the Intergovernmental Panel on Climate Change. Cambridge University Press, Cambridge, United Kingdom and New York, NY, USA, pp. 553–672.

Lesjak, J. and Calderini, D.F., 2017. Increased Night Temperature Negatively Affects Grain Yield, Biomass and Grain Number in Chilean Quinoa. Front Plant Sci, 8. 10.3389/fpls.2017.00352

Liu, X. et al., 2022. The impact of drought and heat stress at flowering on maize kernel filling: Insights from the field and laboratory. Agr Forest Meteorol, 312: 108733. 10.1016/j.agrformet.2021.108733

Lizana, X.C. and Calderini, D.F., 2013. Yield and grain quality of wheat in response to increased temperatures at key periods for grain number and grain weight determination: considerations for the climatic change scenarios of Chile. J Agr Sci, 151(2): 209–221. 10.1017/S0021859612000639

Massuia de Almeida, L.M. et al., 2021. High temperature patterns at the onset of seed maturation determine seed yield and quality in oilseed rape (Brassica napus L.) in relation to sulphur nutrition. Environ Exp Bot, 185: 104400. 10.1016/j.envexpbot.2021.104400

Mayer, L.I., Rattalino Edreira, J.I. and Maddonni, G.A., 2014. Oil Yield Components of Maize Crops Exposed to Heat Stress during Early and Late Grain-Filling Stages. Crop Sci, 54(5): 2236–2250. 10.2135/cropsci2013.11.0795

Meier, U., 2018. Growth stages of mono-and dicotyledonous plants, BBCH Monograph. Julius Kühn-Institut, Quedlinburg, Germany, pp. 204.

Menendez, Y.C., Botto, J.F., Gomez, N.V., Miralles, D.J. and Rondanini, D.P., 2019. Physiological maturity as a function of seed and pod water concentration in spring rapeseed (Brassica napus L.). Field Crop Res, 231: 1–9. 10.1016/j.fcr.2018.11.002

Mera, M., Lizana, X.C. and Calderini, D.F., 2015. Chapter 6 - Cropping systems in environments with high yield potential of southern Chile. In: V.O. Sadras and D.F. Calderini (Editors), Crop Physiology (Second Edition). Academic Press, San Diego, pp. 111–140.

Merrill, A.L. and Watt, B.K., 1973. Energy value of foods: basis and derivation. United States Dept of Agriculture Agriculture handbook. Human Nutrition Research Branch, Agricultural Research Service; for sale by the Supt. of Docs., U.S. Govt. Print. Off., Washington, 105 pp.

Morrison, M.J., 1993. Heat stress during reproduction in summer rape. Can J Botany, 71(2): 303–308. 10.1139/b93-031

Motulsky, H.J., 2014. GraphPad Curve Fitting Guide.

Pokharel, M., Stamm, M., Hein, N.T. and Jagadish, K.S.V., 2021. Heat stress affects floral morphology, silique set and seed quality in chamber and field grown winter canola. J Agron Crop Sci, 207(3): 465–480. 10.1111/jac.12481

Porter, J.R. and Semenov, M.A., 2005. Crop responses to climatic variation. Philosophical Transactions of the Royal Society B: Biological Sciences, 360(1463): 2021–2035. 10.1098/rstb.2005.1752

Poudel, S. et al., 2024. Final Seed Size in Soybean Is Determined during Mid-Seed Filling Stage. Agronomy, 14(4): 763. 10.3390/agronomy14040763

Rao, G.U., Jain, A. and Shivanna, K.R., 1992. Effects of High Temperature Stress on Brassica Pollen: Viability, Germination and Ability to Set Fruits and Seeds. Ann Bot-London, 69(3): 193–198. 10.1093/oxfordjournals.aob.a088329

Rattalino Edreira, J.I., Budakli Carpici, E., Sammarro, D. and Otegui, M.E., 2011. Heat stress effects around flowering on kernel set of temperate and tropical maize hybrids. Field Crop Res, 123(2): 62–73. 10.1016/j.fcr.2011.04.015

Ray, D.K. et al., 2019. Climate change has likely already affected global food production. Plos One, 14(5): e0217148. 10.1371/journal.pone.0217148

Rivelli, G., Calderini, D.F., Abeledo, L.G., Miralles, D.J. and Rondanini, D.P., 2024. Yield and Quality Traits of Wheat and Rapeseed in Response to Source-Sink Ratio and Heat Stress in Post-Flowering. Eur J Agron, 152: 127028. 10.1016/j.eja.2023.127028

Rivelli, G.M. et al., 2021. Assessment of heat stress and cloudiness probabilities in post-flowering of spring wheat and canola in the Southern Cone of South America. Theoretical and Applied Climatology, 145(3-4): 1485–1502. 10.1007/s00704-021-03694-x

Rivelli, G.M. et al., 2023. Photothermal Quotient Describes the Combined Effects of Heat and Shade Stresses on Canola Seed Productivity. Seeds, 2(1): 149–164. 10.3390/seeds2010012

Rodriguez, D. and Sadras, V.O., 2007. The limit to wheat water-use efficiency in eastern Australia. I. Gradients in the radiation environment and atmospheric demand. Aust J Agr Res, 58(4): 287-302. 10.1071/ar06135

Rondanini, D., Mantese, A., Savin, R. and Hall, A.J., 2006. Responses of sunflower yield and grain quality to alternating day/night high temperature regimes during grain filling: Effects of timing, duration and intensity of exposure to stress. Field Crop Res, 96(1): 48–62. 10.1016/j.fcr.2005.05.006

Rondanini, D., Vilariño, M., E. Roberts, M., Polosa, M. and Botto, J., 2014. Physiological responses of spring rapeseed (Brassica napus) to red/far-red ratios and irradiance during pre and post flowering stages, 152. 10.1111/ppl.12227

Sadras, V. and Calderini, D., 2021. Crop Physiology Case Histories for Major Crops. Academic Press, London, United Kingdom, 778 pp.

Sadras, V.O. et al., 2022. Temperature-Driven Developmental Modulation of Yield Response to Nitrogen in Wheat and Maize. Frontiers in Agronomy, 4. 10.3389/fagro.2022.903340

Sadras, V.O. and Moran, M.A., 2013. Nonlinear effects of elevated temperature on grapevine phenology. Agr Forest Meteorol, 173: 107–115. 10.1016/j.agrformet.2012.10.003

Sadras, V.O., Moran, M.A. and Bonada, M., 2013. Effects of elevated temperature in grapevine. I Berry sensory traits. Aust J Grape Wine R, 19(1): 95–106. 10.1111/ajgw.12007

Sandaña, P.A., Harcha, C.I. and Calderini, D.F., 2009. Sensitivity of yield and grain nitrogen concentration of wheat, lupin and pea to source reduction during grain filling. A comparative survey under high yielding conditions. Field Crop Res, 114(2): 233–243. 10.1016/j.fcr.2009.08.003

Secchi, M.A. et al., 2023. Effects of heat and drought on canola (Brassica napus L.) yield, oil, and protein: A meta-analysis. Field Crop Res, 293: 108848. 10.1016/j.fcr.2023.108848

Sehgal, A. et al., 2017. Effects of drought, heat and their interaction on the growth, yield and photosynthetic function of lentil (Lens culinaris Medikus) genotypes varying in heat and drought sensitivity. Front Plant Sci, 8. 10.3389/fpls.2017.01776

Slafer, G. and Rawson, H., 1994. Sensitivity of wheat phasic development to major environmental factors: a re-examination of some assumptions made by physiologists and modellers. Funct Plant Biol, 21(4): 393–426. 10.1071/PP9940393

Slafer, G.A., Savin, R. and Sadras, V.O., 2014. Coarse and fine regulation of wheat yield components in response to genotype and environment. Field Crop Res, 157: 71–83. 10.1016/j.fcr.2013.12.004

Soba, D., Arrese-Igor, C. and Aranjuelo, I., 2022. Additive effects of heatwave and water stresses on soybean seed yield is caused by impaired carbon assimilation at pod formation but not at flowering. Plant Sci, 321: 111320. 10.1016/j.plantsci.2022.111320

Stone, P. and Nicolas, M., 1995. Effect of timing of heat stress during grain filling on two wheat varieties differing in heat tolerance. I. Grain growth. Funct Plant Biol, 22(6): 927–934. 10.1071/PP9950927

Triboi-Blondel, A.-M. and Renard, M., 1999. Effects of temperature and water stress on fatty acid composition of rapeseed oil, Proceedings of the 10th Intetrnational Rapeseed Congress. CGIRC, Canberra, Australia.

Trnka, M., Dubrovský, M. and Žalud, Z., 2004. Climate change impacts and adaptation strategies in spring barley production in the Czech Republic. Climatic Change, 64(1): 227–255. 10.1023/B:CLIM.0000024675.39030.96

UFOP, 2024. Report on Global Market Supply 2023/2024, Union zur Förderung von Oel-und Proteinpflanzen e. V., Berlin, Germany.

Varshney, R.K. et al., 2021. Designing future crops: genomics-assisted breeding comes of age. Trends Plant Sci, 26(6): 631–649. 10.1016/j.tplants.2021.03.010

Verdejo, J. and Calderini, D.F., 2020. Plasticity of seed weight in winter and spring rapeseed is higher in a narrow but different window after flowering. Field Crop Res, 250: 107777. 10.1016/j.fcr.2020.107777

Wang, N. et al., 2022. Impacts of heat stress around flowering on growth and development dynamic of maize (Zea mays L.) ear and yield formation. Plants (Basel, Switzerland), 11(24). 10.3390/plants11243515

Wardlaw, I.F., Blumenthal, C., Larroque, O. and Wrigley, C.W., 2002. Contrasting effects of chronic heat stress and heat shock on kernel weight and flour quality in wheat. Funct Plant Biol, 29(1): 25–34. 10.1071/PP00147

Zhang, Z. et al., 2022. Effects of post-anthesis temperature and radiation on grain filling and protein quality of wheat (Triticum aestivum L.). Agronomy, 12(11). 10.3390/agronomy12112617

Zheng, M., Terzaghi, W., Wang, H. and Hua, W., 2022. Integrated strategies for increasing rapeseed yield. Trends Plant Sci, 27(8): 742–745. 10.1016/j.tplants.2022.03.008

Zhu, Y. et al., 2012. Analysis of gene expression profiles of two near-isogenic lines differing at a QTL region affecting oil content at high temperatures during seed maturation in oilseed rape (Brassica napus L.). Theor Appl Genet, 124(3): 515–531. 10.1007/s00122-011-1725-2

